# A harmonized functional reference framework for *GBA1*

**DOI:** 10.64898/2026.11.15.766795

**Authors:** Sophie Glendinning, Vahid Aslanzadeh, Marguerite Clarke, Yustika Sari, Karolina Zeneviciute, Pilar Alvarez Jerez, Lara M. Lange, Cornelis Blauwendraat, Global Parkinson’s Genetics Program (GP2), Pawel Lis, Katja Lohmann, Christine Klein, Miratul M. Muqit, Dario R. Alessi, Grzegorz Kudla, Esther Sammler

**Affiliations:** MRC Protein Phosphorylation and Ubiquitylation Unit, Faculty of Life Sciences, University of Dundee, Dundee, UK; MRC Human Genetics Unit, Institute of Genetics and Cancer, University of Edinburgh, Crewe Road, EH4 2XU, Edinburgh, UK; Global Parkinson’s Genetic Program (GP2), Chevy Chase, MD, USA; Department of Neurology, University Medical Center Hamburg-Eppendorf, Hamburg, Germany; Coalition for Aligning Science, Chevy Chase, MD, USA; Institute of Neurogenetics, University of Luebeck, Luebeck, Germany; Section of Movement Disorders, Department of Neurology, University of Lübeck, Lübeck, Germany; UK Dementia Research Institute Parkinson’s Research Centre at University of Edinburgh, Edinburgh, UK; Institute for Neuroscience and Cardiovascular Research, School of Neurological and Cardiovascular Sciences, College Medicine and Veterinary Medicine, The University of Edinburgh, UK; Aligning Science Across Parkinson’s (ASAP) Collaborative Research Network, Chevy Chase, MD 20815, USA; Department of Neuroscience, School of Medicine, University of Dundee, Dundee, UK

**Author notes:** Corresponding authors. Correspondence should be addressed to Grzegorz Kudla or Esther Sammler. These authors contributed equally to this work (Sophie Glendinning, Vahid Aslanzadeh).

## Abstract

Systematic functional evidence is lacking for most rare *GBA1* variants, limiting their interpretation in Gaucher disease (GD) and Parkinson’s disease (PD). Here, we used a multiplexed assay of variant effect (MAVE) to generate a comprehensive atlas of lysosome-specific glucocerebrosidase (GCase) activity for more than 10,400 *GBA1* variants, covering 97% of all possible single-amino-acid substitutions. The atlas recapitulated known structural and biochemical determinants of glucocerebrosidase function and revealed concordance between functional impairment and expert-curated pathogenicity classifications. It identified significantly reduced activity in 94 of 186 reported variants of uncertain significance and revealed a quantitative relationship between functional impairment and GD disease severity. Integration with the Global Parkinson’s Genetics Program associated greater variant-level functional impairment with earlier PD onset and higher PD odds ratios. Independent measurements of lysosomal glucocerebrosidase activity in stable cell lines expressing individual variants validated the MAVE results and resolved subtle functional deficits among variants close to the wild-type range, including established PD risk alleles. These findings provide a functional reference framework linking *GBA1* variants to their molecular consequences and clinical phenotypes and demonstrate how comprehensive variant-effect maps can inform genetic interpretation and precision medicine.

## Introduction

*GBA1* encodes the lysosomal enzyme β-glucocerebrosidase (GCase). Within the acidic environment of the lysosome, GCase catalyses the hydrolysis of glucosylceramide (GlcCer) into ceramide and glucose. Gaucher disease (GD), a lysosomal storage disorder, results from biallelic pathogenic variants in *GBA1* that lead to loss of GCase activity (BRADY et al., 1965; Tsuji et al., 1987). Reduced GCase activity results in the accumulation of its substrate, GlcCer and its deacylated derivative glucosylsphingosine (GlcSph), and has been implicated in the accumulation and aggregation of α-synuclein (Dekker et al., 2011; Gegg et al., 2022). In addition, large cohort studies and genome-wide association studies (GWAS) have identified genetic variation in *GBA1* as one of the most common genetic risk factors for Parkinson disease (PD) (Nalls et al., 2014, 2019; Sidransky et al., 2009). Risk-associated coding variants in *GBA1* are present in approximately 5-15% of individuals with PD, although prevalence varies across populations (Vieira et al., 2024). Notably, an intronic *GBA1* risk variant has even been reported in up to 50% of African individuals with PD (Rizig et al., 2023). This variant has also been shown to reduce GCase activity in a dosage-dependent manner through altered splicing (Álvarez Jerez et al., 2024). The frequency of coding risk-associated variants is particularly high among individuals of Ashkenazi Jewish ancestry, where *GBA1* variants, predominantly N409S, have been reported in up to 20% of patients with PD (Vieira et al., 2024). Despite this strong genetic link, PD penetrance among *GBA1* variant carriers is incomplete, and the majority of variant carriers will not develop PD during their lifetime. In addition to increased disease risk, PD in *GBA1* variant carriers is associated with prodromal REM sleep behavior disorder and a more aggressive clinical phenotype, including earlier age at onset, faster motor progression and an increased risk of cognitive impairment (Gan-Or et al., 2015; Hassanin et al., 2025; Malek et al., 2018; Menozzi et al., 2025; Zhao et al., 2016). As multiple *GBA1-*targeted therapeutic approaches - including substrate reduction therapies, pharmacological chaperones, allosteric activators and gene therapies - advance through clinical development, accurate interpretation of *GBA1* variants is becoming increasingly important for patient stratification, clinical trial enrolment and future precision medicine.

The widespread implementation of next-generation sequencing has revealed extensive genetic variation within *GBA1*, including missense, nonsense, splice-site and regulatory variants, together with complex structural rearrangements resulting from recombination with the highly homologous *GBA1LP* pseudogene (Rossi et al., 2025). Although structural and non-coding variants contribute substantially to disease, missense variants represent the largest class encountered (Rossi et al., 2025). While recurrent variants such as N409S and L483P have been extensively characterised, most missense variants have been reported in only a small number of individuals or single families, leaving their clinical significance uncertain.

Interpretation of *GBA1* variants is further complicated because different alleles contribute to disease risk through distinct biological mechanisms and clinical consequences (Chatterjee & Krainc, 2023). Furthermore, variants such as E365K and T408M increase PD risk without being established causes of GD. Consequently, while ACMG/AMP guidelines provide an appropriate framework for classifying pathogenic variants in recessive GD, they are less readily applicable to PD, leaving the clinical significance of many rare *GBA1* variants uncertain (Badonyi & Marsh, 2025; Rossi et al., 2025). In the context of PD, variant interpretation relies on case-control association studies, functional evidence and biological plausibility rather than classic monogenic criteria. For novel or extremely rare variants identified in individual families or singleton cases, interpretation is particularly challenging because sufficient genetic association data and robust functional evidence are often lacking.

Many *GBA1* variants impair protein function through diverse mechanisms, including defective protein folding, reduced stability, impaired lysosomal trafficking and reduced catalytic activity (Stirnemann et al., 2017). Functional assessment is therefore particularly informative for understanding the molecular consequences of individual variants and complements genetic and clinical evidence for variant interpretation. Functional studies have provided important biological insights into a limited number of recurrent *GBA1* variants, including L483P, N409S and E365K (Alcalay et al., 2020; Davis et al., 2025; Galvagnion et al., 2022; Schöndorf et al., 2014; S. Yang et al., 2020; S.-Y. Yang et al., 2017). However, most previous studies have focused on selected variants using relatively low-throughput approaches, limiting their ability to keep pace with the rapidly expanding spectrum of rare variants identified through modern large-scale sequencing. Consequently, comprehensive experimental data remains unavailable for the vast majority of possible coding variants.

Multiplex assays of variant effect (MAVEs) have emerged as a powerful approach for systematic functional annotation of human genetic variation (McEwen et al., 2026). By enabling the parallel assessment of thousands of variants within a single experiment, MAVEs generate comprehensive functional landscapes that complement genetic and clinical evidence for variant interpretation. Here, we generated a comprehensive functional atlas of *GBA1* consisting of 10,425 experimentally characterized coding variants. By integrating multiplex functional profiling with clinical annotation and validation in individual cell lines, we sought to establish a harmonised functional reference framework for the interpretation of *GBA1* variation in GD and PD. This resource provides functional evidence for variant interpretation at scale and establishes a framework for improving variant classification, genetic diagnosis, and precision medicine in both diseases. Although this approach does not capture pathogenic structural rearrangements involving *GBA1LP* or non-coding regulatory variants, it represents an important step towards comprehensive functional annotation of *GBA1*.

## Results

### Development of a multiplexed lysosomal GCase activity assay for systematic variant effect mapping

To enable systematic functional interrogation of *GBA1* coding variation, we developed a fluorescence-activated cell sorting (FACS) assay to quantify lysosomal GCase activity in living cells. The assay utilises LysoFix-GBA, a cell-permeable fluorogenic substrate that selectively accumulates within lysosomes (Zhu et al., 2023). Following hydrolysis by active GCase, the quencher moiety is cleaved, generating a fluorescent signal proportional to lysosomal GCase activity (Fig. 1A). Because fluorescence is generated only following hydrolysis by active lysosomal GCase, the assay provides a cumulative readout of protein expression, folding, intracellular trafficking, lysosomal delivery and catalytic activity.

**Figure 1:**
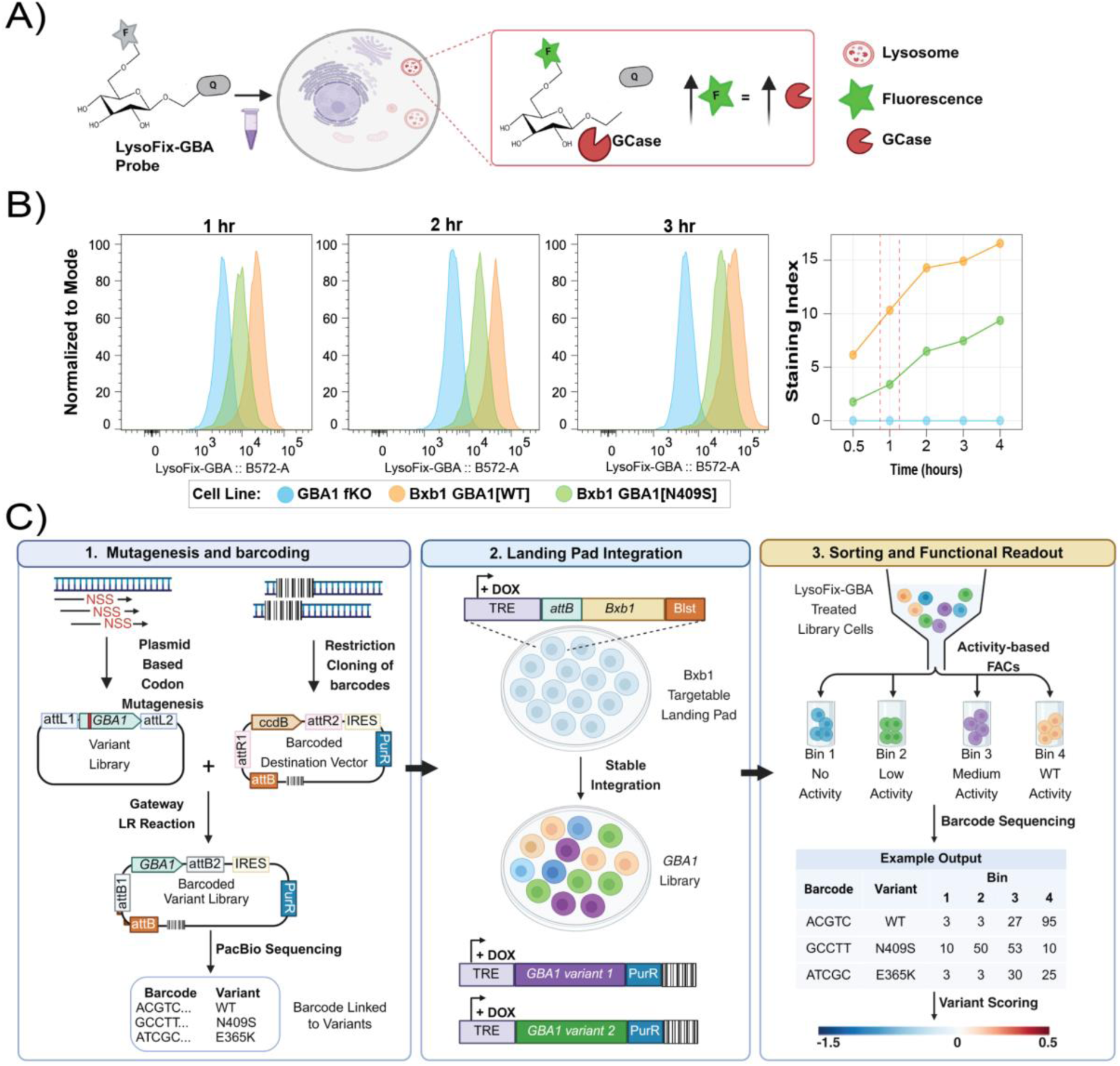
Development of a multiplexed assay for functional assessment of *GBA1* variants. A) Schematic of the LysoFix-GBA assay. The fluorescently quenched LysoFix-GBA substrate is internalized by live cells and hydrolyzed by lysosomal GCase releasing fluorescence proportional to GCase enzymatic activity. B) Optimization of LysoFix-GBA incubation time. *GBA1* fKO HEK Bxb1 landing pad cells (blue), and *GBA1* fKO HEK Bxb1 landing pad cells expressing *GBA1*[WT] (orange), or *GBA1*[N409S] (green) were incubated with 15 μM LysoFix-GBA for 0.5-4 h. Left, representative flow cytometry histograms following LysoFix-GBA incubation for 1hr, 2hrs or 3hrs. Right, time course of LysoFix-GBA signal evaluated using the staining index, the median fluorescent intensity (MFI) normalized to the MFI of *GBA1* fKO / 2* robust SD of *GBA1* fKO. C) Overview of the *GBA1* MAVE workflow. 1. A barcoded saturation library was generated. 2. The library was integrated into the *GBA1* fKO HEK Bxb1 landing-pad cells. 3. The *GBA1* cellular library was sorted by flow cytometry into four fluorescence bins according to GCase activity. Variants within each bin were identified by next-generation sequencing of their barcodes, and the distribution of each variant across bins was used to calculate a functional score.

To establish assay conditions suitable for pooled mutational screening, we first generated a functional *GBA1* knockout (fKO) in HEK293T Bxb1 landing-pad cells (Shukla et al., 2021) (Supplementary Fig. 1), followed by stable single-copy integration of either wild-type (WT) *GBA1* or the pathogenic N409S variant. As expected, knockout cells exhibited minimal LysoFix-GBA fluorescence, whereas WT cells showed a progressive increase in fluorescence over time (Fig. 1B). Cells expressing N409S consistently displayed reduced fluorescence compared with WT, while remaining clearly distinguishable from the knockout, consistent with the reduced GCase activity previously reported for this variant (Fig. 1B) (Alcalay et al., 2015; Sanchez-Martinez et al., 2016; Schöndorf et al., 2014; S. Yang et al., 2020). A one-hour incubation provided robust separation between WT and N409S while maintaining measurements within the linear range of the assay and was therefore selected for subsequent screening. These conditions provided a robust dynamic range suitable for quantitative discrimination between variants with different levels of GCase activity.

A barcoded saturation mutagenesis plasmid library comprising 10,444 coding variants was generated (see Methods), enabling stable single-copy integration of individual *GBA1* variants into HEK-Bxb1 landing pad cells (Fig. 1C). Library composition, variant representation and barcode coverage are summarised in Supplementary Table 1, and Supplementary Fig. 2. Library cells were incubated with LysoFix-GBA for one hour before fixation and sorted by FACS into four bins representing increasing levels of lysosomal GCase activity (Fig. 1C). Genomic DNA was subsequently extracted from each bin, variant-associated barcodes were sequenced, and functional activity scores were calculated from the distribution of each barcode across the four fluorescence bins (Aslanzadeh et al., 2026). Together, these experiments establish a robust multiplexed platform for systematic quantitative functional profiling of *GBA1* variants at scale.

### A comprehensive functional atlas of *GBA1* coding variants

Although several hundred *GBA1* coding variants have been reported in patients to date (Rossi et al., 2025), more than 10,000 non-synonymous single amino acid substitutions are theoretically possible across the *GBA1* coding sequence with its 536 amino acids, excluding the start codon. To systematically define the functional consequences of this coding variant landscape, we generated a comprehensive functional atlas of *GBA1*, experimentally characterising 10,425 of the 10,740 theoretically possible non-synonymous missense and nonsense variants (97.1 % coverage) (Fig. 2A, Supplementary Table 2). The complete dataset is freely available through MaveDB (urn:mavedb:00001327-a) (Rubin et al., 2025) and an interactive web browser (https://gba1-app.onrender.com/). As expected, synonymous variants were functionally indistinguishable from WT, whereas both missense and nonsense variants, on average, exhibited significantly reduced activity (p = 1 × 10⁻³⁰⁰, Supplementary Fig. 3A).

**Figure 2:**
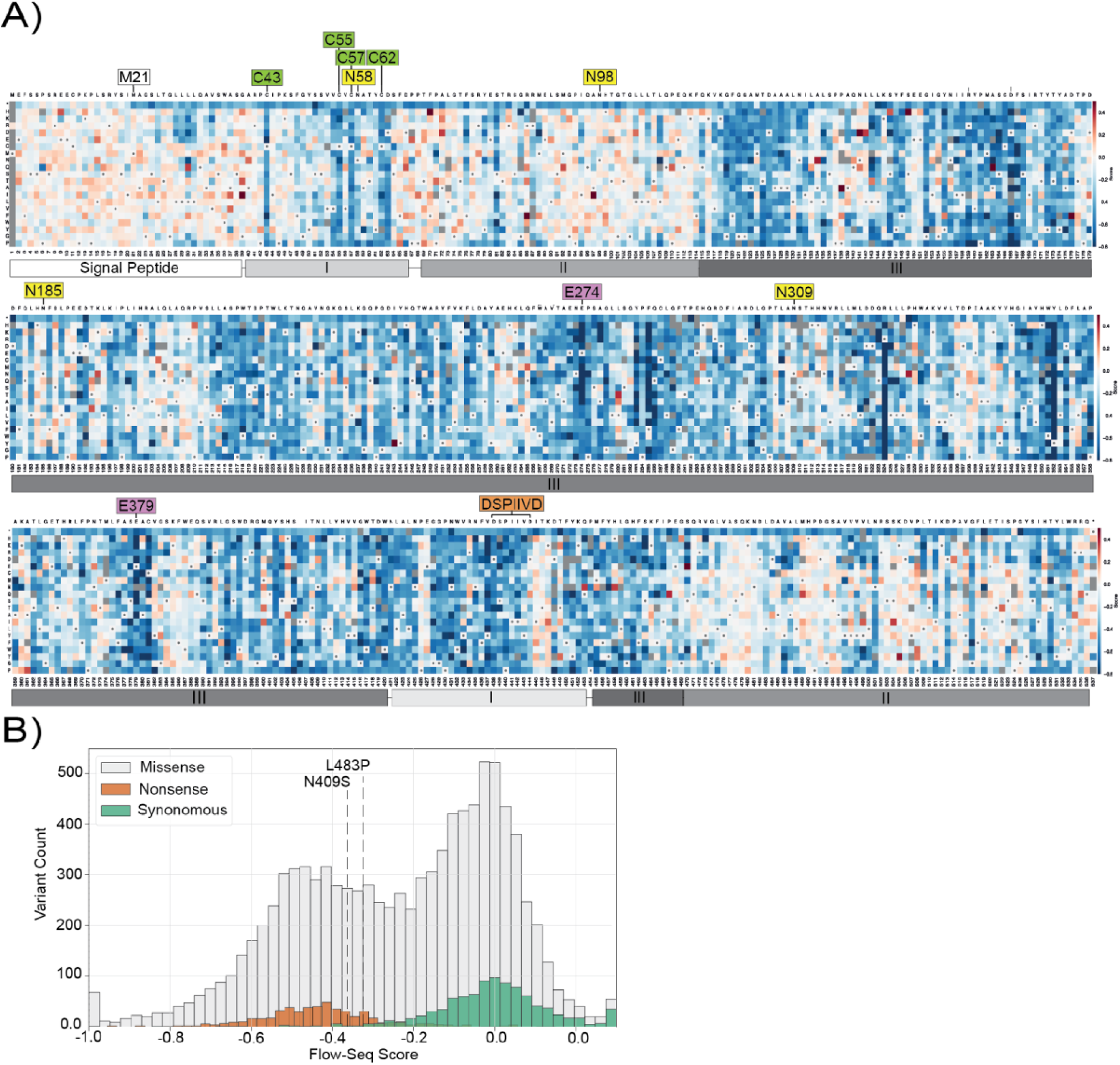
Comprehensive functional atlas of *GBA1* variant effects. A) Heat map showing functional scores for 10,425 *GBA1* missense and nonsense variants measured by multiplex flow-seq following incubation of the variant library with 15 µM LysoFix-GBA for 1 hour. Scores are shown relative to WT *GBA1*, with WT-like variants shown in white, gain-of-function variants in red, and loss-of-function variants in blue. Grey boxes indicate variants for which no functional score could be assigned. *GBA1* domains are indicated below the heat map, including the signal peptide (residues 1-39), domain I (residues 40-66 and 422-453), domain II (residues 69-114 and 470-536), and domain III (residues 115-420 and 455-469). Conserved catalytic residues (purple), N-linked glycosylation sites (yellow), disulphide-forming cysteines (green), the LIMP-2 interactor motif, DSPIIVD (orange), and the M21 start site (white) are highlighted above the protein sequence. B) Distribution of flow-seq scores for missense (grey), nonsense (orange) and synonymous (green) variants. The flow-seq scores of the common PD-associated *GBA1* variants N409S and L483P are indicated.

Inspection of the complete variant effect map revealed distinct patterns of functional constraint across the *GBA1* coding sequence (Fig. 2A). Variants within the N-terminal signal peptide (residues 1-39) generally had little effect on GCase activity, whereas much of the mature protein (residues 40-536) was highly intolerant to amino acid substitutions. Interestingly, nonsense variants upstream of residue 21 retained WT-like activity, consistent with the presence of an alternative translation initiation codon at methionine 21 (Miyoshi et al., 2022; Sorge et al., 1987) (Fig. 2A). Excluding this region, most but not all nonsense variants reduced GCase activity, with a median nonsense flow-seq score of −0.44 (SD = 0.15). To investigate the source of this variability, we used bootstrapping to calculate an empirical p-value for each nonsense variant, testing whether its score differed from wild-type when accounting for barcode coverage (see Methods). Nonsense variants that remained indistinguishable from WT (p ≥ 0.05) had lower barcode coverage than significant ones (median = 2 vs. 8 barcodes; Mann-Whitney U, p = 1.1×10⁻³⁷, Supplementary Figure 3B). Restricting the analysis to variants with ≥3 barcodes increased the fraction of nonsense variants with significantly reduced GCase activity from 81.2% to 91%, while reducing the fraction of total variants retained in the analysis from 97.1% to 84.1% (Supplementary Figure 3C). Increasing the barcode-count threshold beyond 3 further enriched for variants with significantly reduced activity, but progressively reduced the number of nonsense variants retained, highlighting a trade-off between score reliability and library coverage (Supplementary Figure 3C). Among variants with ≥3 barcodes, the 9% of non-significant (p ≥ 0.05) nonsense variants demonstrated low barcode coverage (<7 barcodes) and increased barcode score variability (Supplementary 3B-D).

The variant effect landscape closely reflected the known structural organisation of GCase (Fig. 2A, Supplementary Table 3). Variants in Domain I and the catalytic (β/α)₈ TIM-barrel (Domain III) were generally highly deleterious, whereas the immunoglobulin-like Domain II was comparatively tolerant to missense variation (Supplementary Figure 4). Consistent with established structural biology, residues forming the two Domain I disulfide bridges (Cys43, Cys55, Cys57, and Cys62), which contribute to protein folding and structural stability, were highly intolerant to substitutions (Fig. 2A, Supplementary Figure 5A) (Dvir et al., 2003; Jack et al., 2014; Liou et al., 2006). Similarly, mutations affecting the catalytic residues Glu274 and Glu379 uniformly impaired enzyme activity (Fig. 2A, Supplementary Fig. 5B). Further analysis of residues lining the catalytic cavity revealed that mutations at most positions within this region also resulted in substantial reductions in enzymatic activity, with many exhibiting functional scores lower than those of truncating mutations (Supplementary Fig. 5B) (Romero et al., 2019). In contrast, mutations at residues involved in SapC binding were generally better tolerated, with residue-level flow-seq scores remaining closer to wild-type than those observed for catalytic cavity residues (Supplementary Fig. 5C, F). Mutations at residues within the LIMP-2 binding motif, DSPIIVD (residues 438-444) identified by (Liou et al., 2014) were generally deleterious (Supplementary Fig. 5D).

Similarly, the functional importance of key post-translational modifications was recapitulated by the MAVE. GCase undergoes N-linked glycosylation at residues N58, N98, N185 and N309, which contributes to protein folding and lysosomal trafficking. Variants affecting N58 produced marked reductions in activity, whereas substitutions at N98, N185, and N309 were largely tolerated and indistinguishable from WT (Fig. 2A, Supplementary Fig. 5E). Consistent with previous findings (Berg-Fussman et al., 1993), our data show that glycosylation at N58 is important for the production of catalytically active *GBA1*, whereas mutagenesis of the other occupied glycosylation sites did not substantially affect catalytic activity. The assay also accurately captured the effect of clinically well-established *GBA1* variants. Both N409S and L483P exhibited marked reductions in GCase activity (Fig. 2B). These measurements were supported by robust barcode representation (N409S: 8 barcodes, p = 0.0016; L483P: 28 barcodes, p = 0.0002) and are consistent with numerous previous studies demonstrating reduced GCase activity associated with these variants (Alcalay et al., 2015; Galvagnion et al., 2022; García-Sanz et al., 2017; Sanchez-Martinez et al., 2016; Schöndorf et al., 2014; Thomas et al., 2021; S. Yang et al., 2020).

To further assess the biological relevance of the variant effect map, flow-seq scores were compared with independent measures of evolutionary conservation and computational variant effect prediction (VEP). Functional scores showed a strong inverse correlation with residue conservation determined by ConSurf (https://consurf.tau.ac.il/) (Spearman’s ρ = −0.72, Supplementary Fig. 6), indicating that substitutions at evolutionarily conserved residues were associated with greater functional impairment. Flow-seq scores were also concordant with computational VEPs, including AlphaMissense (Spearman’s ρ = −0.733), CADD (Spearman’s ρ = −0.622) and REVEL (Spearman’s ρ = −0.683), among others (Supplementary Fig. 7, Supplementary Table 4), demonstrating strong agreement between experimental measurements and state-of-the-art computational predictions. Consistent with these findings, flow-seq scores also correlated strongly with the first principal component reported by (Lanore et al., 2025), which integrates structural, evolutionary, and clinical features of *GBA1* variation (Supplementary Table 4).

Together, these data demonstrate that multiplex functional profiling accurately captures the structural and biochemical constraints governing GCase function while providing systematic experimental annotation for almost all possible *GBA1* coding variants.

### Multiplex functional profiling complements clinical interpretation of *GBA1* variation

To evaluate the clinical relevance of the functional atlas, we compared flow-seq scores with expertly curated *GBA1* variant annotations from the International Parkinson and Movement Disorder Society Genetic mutation database (https://www.mdsgene.org). A recent systematic review by MDSGene identified 761 unique *GBA1* codon variants reported across 27,963 individuals with PD, GD, GD with PD, or other parkinsonian disorders (Rossi et al., 2025). Of these, 494 were missense or nonsense codon variants, while the remaining variants comprised a range of other variant types, including splice-affecting variants, indels, and complex rearrangements (Supplementary Fig 8A). At the protein level, these 494 missense and nonsense codon variants represented 486 unique protein coding variants, of which 477 (98%) were represented in our functional atlas.

Flow-seq scores closely recapitulated established ACMG/AMP classifications used for monogenic disorders (Fig. 3A). Variants classified as pathogenic or likely pathogenic exhibited marked reductions in GCase activity, with 82 of 111 pathogenic variants (73.9%) and 130 of 172 likely pathogenic variants (75.6%) demonstrating significantly reduced activity (*p* < 0.05) relative to WT, and sufficient barcode coverage (≥ 3) (Fig 3A, Supplementary Fig. 9A). Many of the remaining pathogenic or likely pathogenic variants demonstrated low barcode coverage and/or a moderate reduction in activity which did not meet the statistical threshold. It is therefore possible that milder effects on activity are below the sensitivity limit for statistical significance within this large-scale activity screen. In contrast, variants classified as likely benign were generally not significantly different from WT in the multiplexed assay (Supplementary Fig. 9A). The variant E365K, classified as benign with regards to GD following ACMG/AMP but known to be a PD risk factor, was represented by only a single barcode and was therefore considered to have insufficient confidence for functional interpretation in the multiplexed assay.

**Figure 3:**
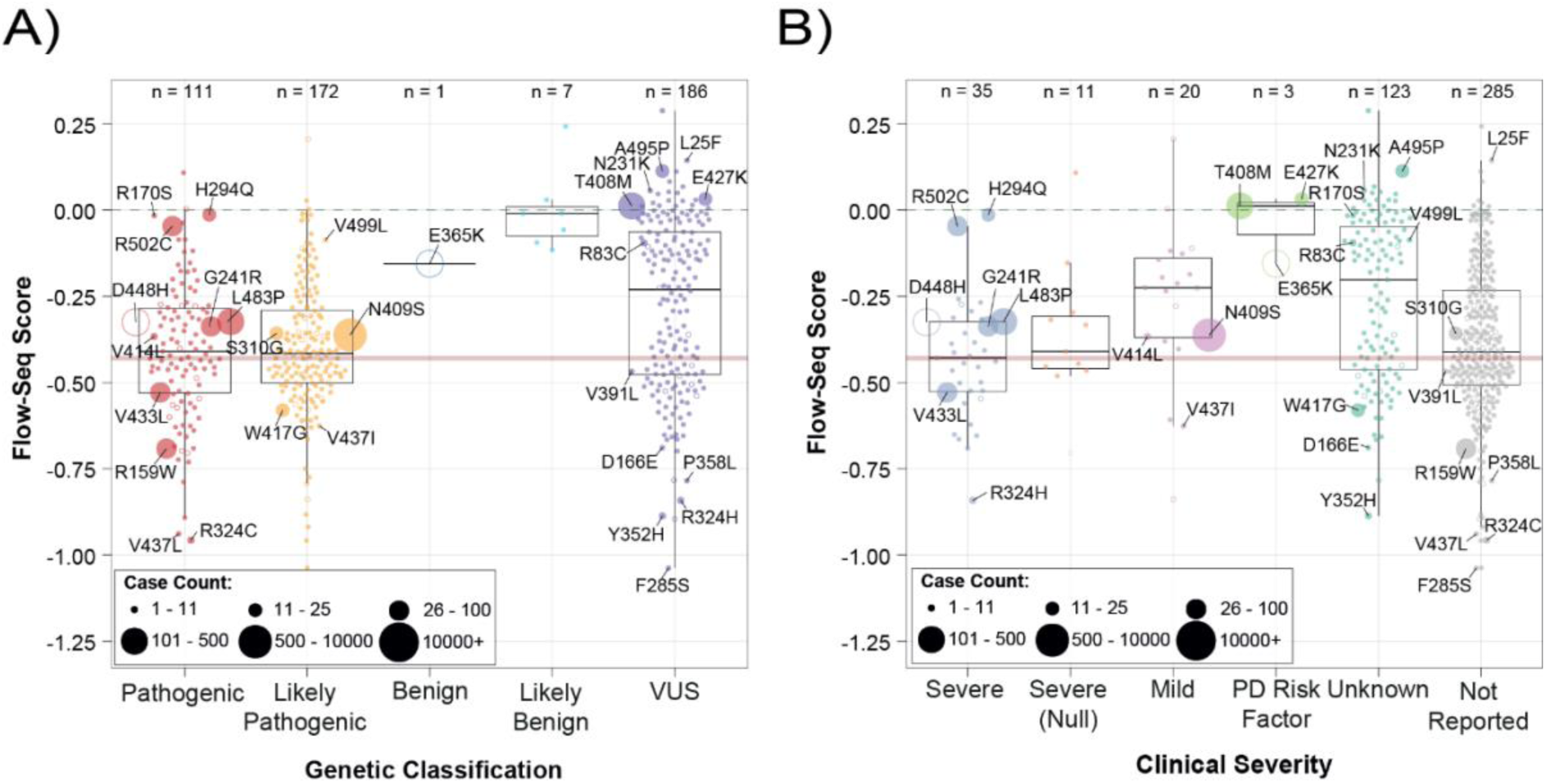
Relationship between experimentally derived *GBA1* functional flow-seq scores and clinical variant annotations. Each circle represents a unique *GBA1* variant reported in MDSGene, with circle size proportional to the number of individuals carrying each variant. Filled and open circles denote variants with ≥ 3 barcodes or < 3 barcodes, respectively. A) Functional flow-seq scores of *GBA1* variants grouped according to their pathogenicity classification: pathogenic (n = 111), likely pathogenic (n = 172), benign (n = 1), likely benign (n = 7), or VUS (n = 186). B) Functional flow-seq scores of *GBA1* variants grouped according to their clinical severity annotation: severe (n = 35), severe null (n = 11), mild (n = 20), PD risk factor (n = 3), unknown (n =123) or unannotated (n = 285).

Variants classified as variants of uncertain significance (VUS) spanned the full spectrum of functional activity (Fig. 3A). Of the 186 VUS included in the analysis with sufficient barcode coverage (≥ 3), 94 (50.5%) showed significantly reduced activity relative to WT (*p* < 0.05), whereas 82 (44.1%) were not significantly different from WT (*p* > 0.05); and a further 10 (5.4%) had insufficient barcode coverage for functional interpretation (< 3 barcodes) (Fig. 3A, Supplementary Fig. 9B). These findings demonstrate that multiplex functional profiling revealed biological differences among clinically unresolved *GBA1* variants, identifying a substantial subset with marked reductions in lysosomal GCase activity.

Although ACMG/AMP classification provides an established framework for interpreting *GBA1* variants in GD, it is not designed to capture PD risk variants, i.e., E365K and T408M. For example, E365K and T408M are classified as benign and as a VUS, respectively, under the ACMG guideline-based MDSGene classification framework, despite both being established PD risk factors (Rossi et al., 2025). We therefore assessed the relationship between flow-seq scores and variants classified as PD risk factors, alongside variants associated with mild and severe GD phenotypes. Using clinical severity classifications from the GBA1-PD Browser (Parlar et al., 2023), as reported in MDSGene (Rossi et al., 2025), functional scores closely tracked the established clinical spectrum of GD severity (Fig. 3B; Supplementary Fig. 8B). Variants associated with severe or severe (null) GD (mean score −0.411) scored significantly lower than variants associated with mild GD (mean score −0.271; BH-adjusted p = 0.0227) or PD risk variants (mean score −0.037; BH-adjusted p = 0.0175, Fig. 3B). In contrast, PD risk variants clustered close to WT, potentially reflecting their more subtle functional effects and the greater biological heterogeneity underlying PD susceptibility. Overall, the multiplexed assay captured the broader gradient underlying established clinical severity classifications.

To investigate whether experimentally measured GCase activity was associated with clinical phenotype, we integrated MAVE functional scores with genetic and phenotypic data from the Global Parkinson’s Genetics Program (GP2) (The Global Parkinson’s Genetics Program, 2021). Analysis was restricted to individuals with PD carrying a single heterozygous *GBA1* variant, using harmonized age at motor symptom onset (AAO) data. Flow-seq scores were analysed as continuous variables and compared with variant-level median AAO for variants represented by at least five variant carriers with available data among the European-ancestry cohort. Variants with lower Flow-seq scores were associated with earlier median AAO (Fig. 4A-B, Supplementary Table 5). Among variants with ≥5 carriers, the correlation was Spearman’s ρ = 0.728 (n = 18, Fig 4A), compared with Spearman’s ρ = 0.561 for variants with ≥10 carriers (n = 10, Fig. 4B). Flow-seq scores were also compared with PD odds ratios (ORs) for variants with minor allele counts (MAC) of ≥5 or ≥10. A mild inverse correlation was observed between Flow-seq scores and PD ORs, with variants exhibiting greater functional impairment generally associated with higher PD risk (≥5 MAC: n = 17, Spearman’s ρ = −0.534; Fig 4C; and ≥10 MAC: n = 13, Spearman’s ρ = −0.473; Fig 4D). Overall, the functional atlas showed strong concordance with established GD clinical classifications while providing direct experimental evidence to further stratify clinically unresolved *GBA1* variants. Integration with GP2 data additionally revealed associations between experimentally measured lysosomal GCase activity, PD risk and AAO.

**Figure 4:**
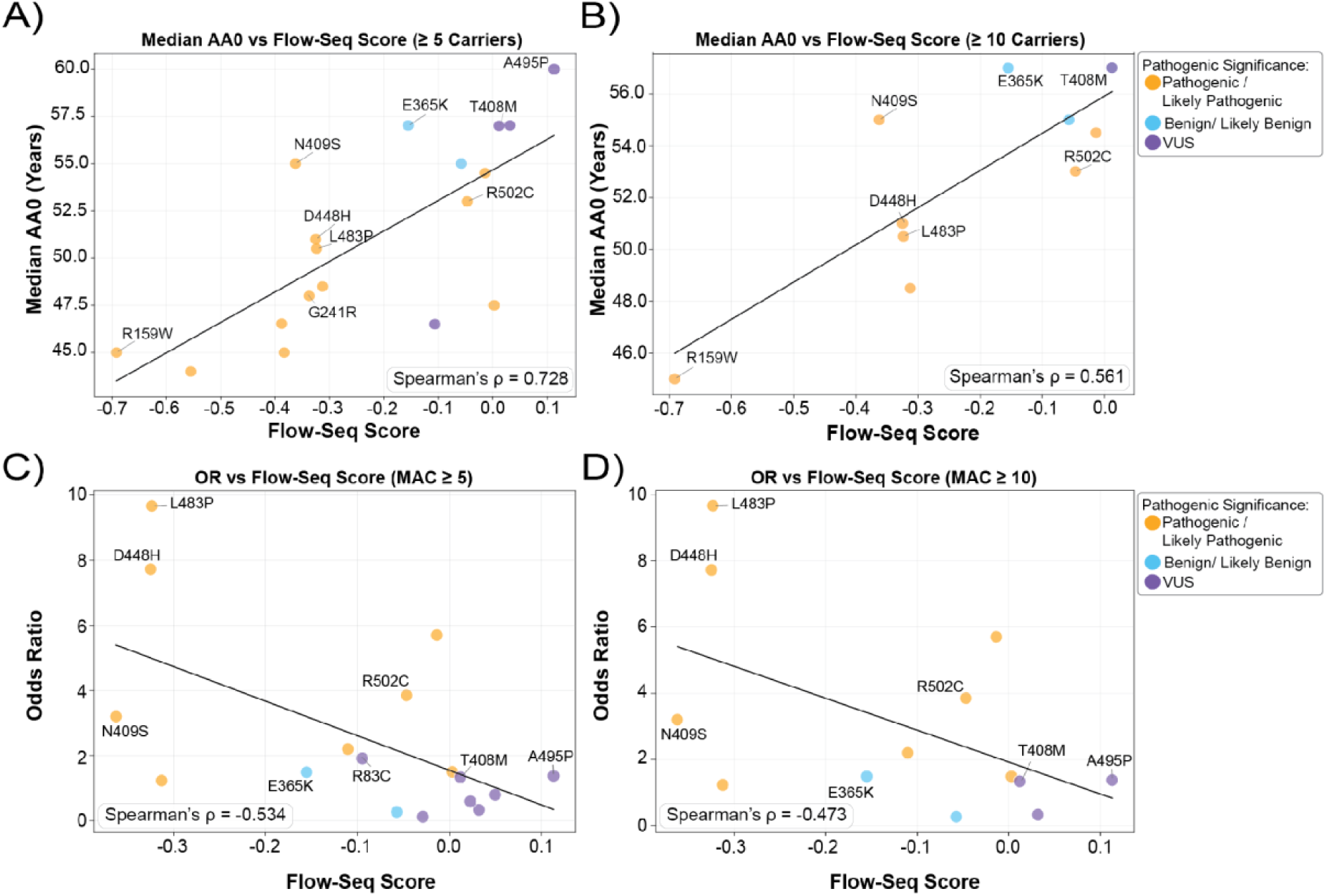
Association of functional scores with Parkinson’s disease age of onset and odds ratios. A-B) Functional scores plotted against the median age of PD onset for each variant, calculated using the GP2 dataset among carriers, with a minimum carrier count of ≥5 (n = 18), or ≥10 (n = 10), respectively. C-D) Functional scores plotted against variant-specific odds ratios (OR) for PD, calculated using the GP2 dataset for variants with a minor allele count (MAC) of ≥5 (n = 17) or ≥10 (n = 13), respectively, and at least one carrier in cases and one carrier in controls. Annotated variants were among those with the highest number of reported cases on MDSGene, and are coloured according to MDSGene genetic annotation: pathogenic/likely pathogenic (orange), benign/likely benign (blue), and VUS (purple).

### Independent validation confirms the accuracy and resolution of the multiplex functional atlas

To independently evaluate the performance of the multiplex assay, we generated stable cell lines expressing individually selected *GBA1* variants spanning the full range of functional effects of the pooled screen. These included severe and mild GD variants, PD risk variants, VUS and variants close to the WT functional range, allowing us to assess both the accuracy and quantitative resolution of the multiplex assay (Table 1, Supplementary Fig. 10). Overall, single-variant measurements showed strong agreement with multiplex flow-seq scores across the full range of GCase activity (Pearson’s *r* = 0.784; Fig. 5C), confirming that pooled functional profiling accurately captures the functional consequences of *GBA1* variation.

**Figure 5:**
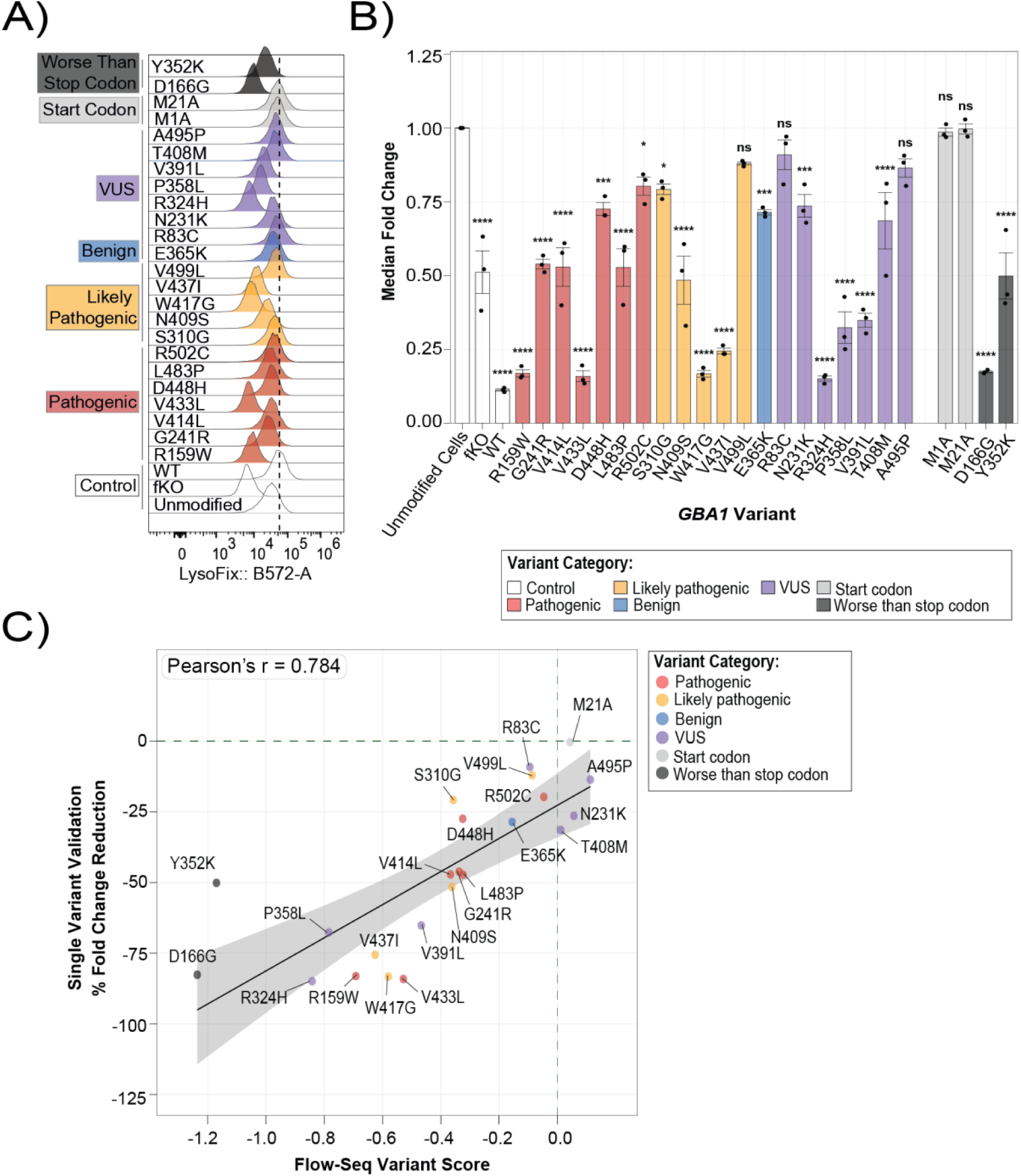
Independent validation of multiplex functional scores. Single-variant HEK Bxb1 landing-pad cell lines representing variants annotated by MDSGene as pathogenic, likely pathogenic, benign and VUS, together with start codon, worse-than-stop-codon, and control lines. Pathogenic *GBA1* variants R159W, G241R, V414L, V433L, D448H, L483P and R502C; likely pathogenic *GBA1* variants S310G, N409S, W417G, V437I and V499L; the benign *GBA1* variant E365K; *GBA1* VUS R83C, T408M, N231K, R324H, P358L, V391L and A495P; start codon variants M1A and M21A; worse-than-stop-codon variants D166G and Y352K; and control lines WT GBA1, *GBA1* fKO cells, and unmodified cells. All cell lines were incubated with 15 μM LysoFix-GBA for 1 hour before flow cytometry analysis. A) Representative flow cytometry histograms showing LysoFix-GBA fluorescence in single-variant lines. B) Quantification of LysoFix-GBA fluorescence expressed as MFI fold change relative to WT *GBA1* from three independent experiments. Data were analyzed using one-way ANOVA with Dunnett’s multiple comparisons test using WT as the reference. Error bars represent mean ± SEM. ns, not significant; *P* < 0.05 (\**), P < 0.01 (**), P < 0.001 (\*\*\**), and *P* < 0.0001 (****). C) Correlation between the MAVE flow-seq functional scores and percentage MFI fold change measured in the corresponding single-variant cell line. The black line indicates the linear regression fit, grey shading represents the 95% confidence interval, and the Pearson correlation coefficient is shown (*r* = 0.784, 95% CI: 0.552–0.904, p = 8.89 × 10⁻⁶).

**Table 1:** Summary of Flow-seq data and MDS gene annotations for variants selected for validation.

| **Variant** | **Flow-Seq Score** | **Flow-Seq Barcode Count** | **Flow-Seq Empirical p Value** | **MDSGene Genetic Classification** | **MDSGene Clinical Severity** | **MDSGene Case Number** |
|---|---|---|---|---|---|---|
| R159W | -0.6917 | 7 | 0.0002 | Pathogenic | - | 212 |
| G241R | -0.3372 | 22 | 0.0002 | Pathogenic | Severe | 150 |
| V414L | -0.3666 | 20 | 0.0002 | Pathogenic | Mild | 15 |
| V433L | -0.5284 | 9 | 0.0002 | Pathogenic | Severe | 148 |
| D448H | -0.3250 | 2 | 0.1446 | Pathogenic | Severe | 530 |
| L483P | -0.3235 | 28 | 0.0002 | Pathogenic | Severe | 6812 |
| R502C | -0.0467 | 9 | 0.6090 | Pathogenic | Severe | 293 |
| S310G | -0.3574 | 5 | 0.01633 | Likely Pathogenic | - | 27 |
| N409S | -0.3624 | 8 | 0.0027 | Likely Pathogenic | Mild | 17890 |
| W417G | -0.5801 | 17 | 0.0002 | Likely Pathogenic | Unknown | 34 |
| V437I | -0.6258 | 9 | 0.0002 | Likely Pathogenic | Mild | 8 |
| V499L | -0.0865 | 5 | 0.4939 | Likely Pathogenic | Unknown | 9 |
| E365K | -0.1552 | 1 | 0.4923 | Benign | Risk Variant | 2643 |
| R83C | -0.0947 | 11 | 0.2736 | VUS | Unknown | 14 |
| N231K | 0.0565 | 8 | 0.5293 | VUS | Unknown | 2 |
| R324H | -0.8418 | 11 | 0.0002 | VUS | Severe | 12 |
| P358L | -0.7845 | 26 | 0.0002 | VUS | - | 3 |
| V391L | -0.4672 | 39 | 0.0002 | VUS | - | 6 |
| T408M | 0.0120 | 4 | 0.9213 | VUS | Risk Variant | 1429 |
| A495P | 0.1132 | 10 | 0.1765 | VUS | Unknown | 70 |
| M1A | - | - | - | - | - | - |
| M21A | 0.0437 | 34 | 0.3408 | - | - | - |
| D166G | -1.2361 | 15 | 0.0002 | - | - | - |
| Y352K | -1.1702 | 6 | 0.0002 | - | - | - |

Pathogenic GD-associated variants (R159W, G241R, V414L, V433L and L483P), together with likely pathogenic variants (S310G, N409S, W417G and V437I), exhibited marked reductions in lysosomal GCase activity when analysed individually (Fig. 5A, B). In contrast, the likely pathogenic variant V499L retained activity comparable to WT in both assays. Amongst VUS, R324H and P358L showed pronounced functional impairment, whereas R83C and A495P remained functionally indistinguishable from WT (Fig. 5A, B), confirming the ability of the multiplex assay to distinguish variants with marked functional impairment from those within the WT-like range.

Individual variant measurements revealed small but reproducible reductions in lysosomal GCase activity for N231K, E365K, T408M, D448H and R502C, none of which differed significantly from WT in the multiplex assay (Fig. 5A, B). This may reflect limited barcode representation for E365K and increased barcode-level variability for D448H, whereas N231K, T408M and R502C were sufficiently represented (Supplementary Fig. 10). Notably, D448H and R502C are classified as GD severe pathogenic variants, while N231K and T408M are classified as VUS (Rossi et al., 2025). T408M, however, is an established PD risk variant that is not associated with GD. Thus, WT-like multiplex scores do not exclude subtle functional deficits detectable in individual variant cell lines.

Total GBA1 protein abundance was largely preserved across all variants analysed. Only R502C showed a modest, non-significant reduction in protein abundance, whereas no significant differences were detected for the remaining variants (Supplementary Fig. 11), indicating that the observed functional defects primarily reflect impaired enzyme activity rather than reduced protein expression.

Beyond validating individual variant effects, the multiplex functional atlas also generated novel mechanistic insights into GBA1 biology. We therefore investigated several variants of particular biological interest. Flow-seq analysis indicated that substitutions upstream of methionine 21 had little or no effect on GCase activity, suggesting that M21 may function as an alternative translation initiation site (Fig. 2A). To test this hypothesis, methionine residues at positions 1 and 21 were individually substituted with alanine. Neither M1A nor M21A significantly altered lysosomal GCase activity, consistent with the possibility that translation can initiate from either methionine residue, as reported previously (Miyoshi et al., 2022; Sorge et al., 1987) (Fig. 5A, B). In addition, the multiplex atlas identified several missense variants with functional scores below the median observed for stop variants (Fig. 2A). Individual analysis confirmed that D166G and Y352K, two variants located within the catalytic cavity, almost completely abolished GCase activity (Fig. 5A, B). Structural mapping placed D166 and Y352 within the local interaction network surrounding the catalytic residues E274 and E379 (Supplementary Fig. 12). Y352 is positioned within the flexible active-site loop of *GBA1* and hydrogen-bonding interactions between its side chain and the two catalytic residues, E274 and E379, support the transition of the loop between extended (inactive) and helical (active) conformations (Lieberman, 2011) (Supplementary Fig. 12). Variants at Y352 could therefore disrupt this conformational switch, alter the positioning or chemical environment of the catalytic glutamates, and consequently impair substrate binding and/or enzymatic activity. D166 contributes directly to substrate binding, providing a potential structural basis for the profound effects on GCase activity (Supplementary Fig. 12). The M1A variant showed a modest, non-significant reduction in total GBA1 protein abundance, whereas no differences were observed for the remaining variants (Supplementary Fig. 11). Together, these validation experiments confirm that multiplex functional profiling accurately captures the functional consequences of *GBA1* variation while highlighting the complementary strengths of targeted single-variant measurements for resolving subtle functional effects. Combined with the comprehensive functional atlas presented here, these findings establish a robust and scalable resource for the functional annotation, biological interpretation and clinical assessment of *GBA1* variation.

## Discussion

The rapid expansion of clinical genetic testing has led to the identification of hundreds of *GBA1* coding variants in patients with GD and PD (Rossi et al., 2025). While a small number of recurrent variants have been extensively characterized, the functional consequences of most reported variants remain unknown, limiting consistent interpretation of rare and newly identified variants as genetics becomes increasingly integrated into clinical care and therapeutic development. Here, we address this gap by generating a comprehensive functional atlas comprising 10,425 experimentally characterised *GBA1* variants, representing 97.1% of all possible coding substitutions. By integrating multiplex functional profiling with curated clinical datasets, independent single-variant validation and patient-derived genetic data, we establish a harmonized functional reference framework for *GBA1* variation that supports variant interpretation, patient stratification and future precision medicine.

Beyond supporting functional variant interpretation, the atlas provides biological insights into the structural determinants of GCase function. The strong concordance between multiplex functional scores and known catalytic residues, disulphide bonds, glycosylation sites and evolutionary conservation demonstrates that the assay faithfully captures the major determinants of lysosomal GCase function (Fig. 2). By systematically assessing almost all possible amino acid substitutions, the atlas extends beyond currently recognised human genetic variation. Although many of these substitutions may never occur in patients, they define the sequence-function relationship for lysosomal GCase activity in our system, whereas those accessible by single-nucleotide change provide pre-emptive functional annotation should they be identified in the future.

In addition to recapitulating established biology, the atlas generated unexpected biological insights that were confirmed by independent single-variant validation. For example, the missense variants D166G and Y352K, located in the catalytic cavity, scored below nonsense variants in the multiplex assay and exhibited markedly impaired lysosomal GCase activity, highlighting the critical importance of individual structural or catalytic residues for enzyme function. Such effects may reflect not only loss of catalytic function, but for some missense variants, additional consequences of mutant protein expression, raising the possibility of mechanisms beyond simple loss of function, e.g. overlapping low- and toxic gain-of-function mechanisms in *GBA1*-associated disease (Smith & Schapira, 2022). Similarly, the tolerance of N-terminal truncating variants highlighted alternative translation initiation at M21, which we confirmed by single-variant analysis, and which is consistent with previous evidence that either *GBA1* start codon can generate active, lysosomally localised GCase (Sorge et al., 1987). Together, these findings illustrate how comprehensive saturation mutagenesis can move beyond functional annotation to uncover new aspects of protein biology while generating testable mechanistic hypotheses.

A major advance of this work is that the functional atlas was benchmarked at scale against clinical and genetic data. Although expert curation, segregation studies and population genetics remain the cornerstones of variant classification, systematic functional evidence has historically been available for only a small number of *GBA1* variants. The strong concordance between multiplex functional scores, established GD pathogenicity classifications and recognised disease severity, together with the association of functional scores with PD age at onset and odds ratios, demonstrates the clinical relevance of systematic experimental functional profiling.

Perhaps most importantly, the atlas provides functional evidence to inform the interpretation of VUS. More than half (94 of 186; 50.5%) of clinically unresolved variants demonstrated clear reductions in lysosomal GCase activity, whereas a similarly large proportion (44.1%) fell within the wild-type functional range. Importantly, experimentally demonstrated functional impairment does not, in isolation, establish clinical pathogenicity or PD risk. The interpretation framework was deliberately designed to prioritise specificity, such that variants classified within the wild-type functional range cannot necessarily be considered functionally neutral and may include variants with subtle loss of function that fall below the resolution of the multiplex assay.

Although the atlas readily complements clinical interpretation in GD, its greatest translational potential may lie in PD. Whereas biallelic *GBA1* variants directly cause GD by lysosomal enzyme deficiency, heterozygous variants primarily act as incompletely penetrant risk factors that increase PD disease susceptibility and modify clinical phenotype. Unlike GD, where variant classification is based on established criteria for monogenic disease, PD presents a more nuanced challenge: understanding how quantitative differences in lysosomal GCase function contribute to disease susceptibility and clinical heterogeneity, including subtle functional effects that may increase PD risk without causing GD.

Several *GBA1* variants associated with increased PD risk, including E365K and T408M, showed small, reproducible reductions in lysosomal GCase activity when analysed individually, although T408M was WT-like and E365K had insufficient barcode coverage for interpretation in the multiplex assay. This highlights the complementary sensitivity of the two approaches: the multiplex assay robustly identifies moderate-to-severe functional impairment at scale, whereas individual variant measurements can resolve more subtle effects close to WT. These findings reinforce that WT-like multiplex score do not necessarily exclude subtle functional impairment, particularly for variants associated with PD risk. Functional scores should therefore be interpreted as a continuum and integrated with genetic association, clinical and segregation data rather than used as a standalone classification of PD risk. Additionally, GCase functional activity scores may not fully capture the spectrum of PD-relevant *GBA1* biology, as some variants may influence disease risk through effects on α-synuclein handling, lipid metabolism or cellular stress responses that are not linearly reflected by residual enzyme activity.

Consistent with this, integration of the functional atlas with genetic and phenotypic data from GP2 revealed associations between greater variant-level functional impairment with earlier PD onset and higher PD odds ratios. Although these variant-level associations require confirmation in larger and independent datasets, they suggest that quantitative functional profiling may help capture biologically meaningful variation in *GBA1*-associated PD and inform future patient stratification and genetically targeted clinical trials.

The use of a single experimental system was a deliberate design choice that enabled large-scale comparison of more than 10,000 variants under identical experimental conditions while minimizing the confounding influence of genetic background. A major strength of the multiplex assay is that it provides an integrated measure of lysosomal GCase function, reflecting the combined effects of protein expression, folding, maturation, intracellular trafficking, post-translational modification and catalytic activity. However, this integrated readout does not distinguish the specific molecular mechanism underlying functional impairment, and variants identified through the atlas provide a basis for prioritizing targeted mechanistic studies. The atlas also identified a small number of variants with apparent increases in GCase activity relative to WT; however, these effects were generally modest, showed limited statistical support and did not cluster within a clear structural region. If validated, such variants could provide insights into mechanisms that enhance GCase function and potentially inform therapeutic strategies, but independent validation in orthogonal assays and relevant cellular models will be required to establish their biological significance.

A further limitation is that the assay was performed in an isogenic stable overexpression system in HEK Bxb1 landing pad cells. Although this provides a robust systematic platform for large-scale functional profiling, it does not fully recapitulate endogenous *GBA1* expression, regulation or the heterozygous genetic context present in most individuals with PD. As the assay uses a cDNA-based construct, it also does not capture effects on splicing or complex rearrangements involving the GBA1LP pseudogene. Future work should extend this framework to endogenous genome-edited systems and disease-relevant cell types, including iPSC-derived dopaminergic neurons and microglia, to determine the extent to which variant effects are conserved across biological contexts. Future iterations of multiplex functional profiling may further improve discrimination among variants with near-wild-type activity through optimization of fluorescence binning strategies and other refinements to increase sensitivity for detecting subtle functional effects while maintaining the scalability of the approach.

More broadly, our study illustrates how comprehensive functional atlases can bridge experimental biology, clinical genetics and patient-derived datasets. Together with our recent application of the same approach to *PRKN* (Aslanzadeh et al., 2026) these findings demonstrate the broader applicability of MAVE methodologies to disease genes with scalable functional readouts. Such experimental reference frameworks have the potential to advance rare variant interpretation, biologically informed patient stratification, and precision medicine across neurogenetics.

## Methods

### General experimental reagents

#### Cell Culture

HEK293T cells carrying the G542A_pLenti-Tet-coBxb1-2A-BFP_IRES_iCasp9-2A-Blast_rtTA3 landing pad (gift from K. Matreyek lab) (Shukla et al., 2021) were maintained in DMEM supplemented with 10% fetal calf serum, 2 mM L-glutamine, penicillin (100 U/mL), streptomycin (100 μg/mL). HEK landing pad cells transfected with *GBA1* variants were supplemented with 3 μg/mL doxycycline (Sigma) and 1.5 μg/mL puromycin (Sigma).

#### Generating a GBA1 fKO Cell Line

A paired Cas9-nickase CRISPR/Cas9 strategy was implemented to generate a *GBA1*-functional knockout (*GBA1*-fKO) HEK293T Bxb1 landing pad cell line. The generation of *GBA1* fKO HEK293T Bxb1 landing pad cells, screening, and subsequent sequencing was performed as described in the protocol available on protocols.io (doi: 10.17504/protocols.io.xygxmkp5ol8j/v1). Two guide RNAs were designed to target exon 6. Guide 1 (5’-GAATGGGAAGGGGTCACTCAA-3’) was cloned into the puromycin-selectable pBabe-PU6 plasmid (DU48788, https://mrcppureagents.dundee.ac.uk/), and guide 2 (5’-GCCATTGGTCTTGAGCCAAGT-3’) was cloned into the spCas9-D10A-expressing pX335 vector (Addgene Plasmid #42335), resulting in the constructs (DU69440 and DU 69441, https://mrcppureagents.dundee.ac.uk/). Sequencing results were aligned against the wild-type GBA1 exon 6 sequence using SnapGene v8.0.1. A 45-bp in-frame deletion within exon 6 was confirmed (Zenodo doi: 10.5281/zenodo.23169564), resulting in the absence of *GBA1* protein expression and enzymatic activity (Supplementary Fig 1).

#### LysoFix-GBA GCase activity assay

The LysoFix-GBA substrate was synthesised by WuXi AppTec following the synthetic strategy previously reported by (Zhu et al., 2023). The LysoFix-GBA GCase activity assay was carried out following the published protocol available on protocols.io (doi: 10.17504/protocols.io.4egnxb8mml5d/v1). For the GCase assay, cells were incubated with 15 µM LysoFix-GBA or vehicle control (DMSO) at 37°C in 5% CO₂. Single variant lines were incubated for the indicated time points for the time course experiments, and within 24 hours of preparation, samples were run on the NovoCyte (ACEA Biosciences). In preparation for the GCase activity MAVE, 140 million cells from the *GBA1* library were processed for FACS-based sorting. Library cells were incubated for 1 hour with LysoFix-GBA, and within 24 hours of preparation of samples for flow-cytometry were single-cell sorted using the FACS Aria II (BD Biosciences).

#### Sample Lysis, Preparation, and Immunoblotting

Immunoblotting preparation was performed as described in the protocol available on protocols.io (doi: 10.17504/protocols.io.epv567z3ng1b/v1). Protein bands were acquired via near-infrared fluorescent detection using the Odyssey CLx imaging system and quantified using the Image Studio software.

#### Antibodies

Mouse monoclonal GBA (clone 2E2) (AP1140, Calbiochem) was used at a dilution of 1:1000 and mouse monoclonal anti-GAPDH (sc-32233, Santa Cruz Biotechnology) was used at a dilution of 1:10,000. The secondary antibody, donkey anti-mouse (LI-COR, 926-32212) was used at a 1:20,000 dilution.

#### Plasmid Construction

The human *GBA1* coding sequence was subcloned from pcDNA5D FRT/TO/GBA1[WT] (MRC PPU Reagents and Services, University of Dundee, DU80234) into the entry vector pAINt (pAINt, Addgene_262847) (McDonnell et al., 2024). Plasmid cloning, transformation into competent *E.coli* cells, and plasmid isolation was performed as described in the protocol available on protocols.io (doi: 10.17504/protocols.io.byl42exyovo5/v1). The sequence of the final pAINT-*GBA1* entry plasmid (Addgene_263442) was confirmed with whole plasmid sequencing. Saturation mutagenesis was performed on this pAINT-*GBA1* entry plasmid.

#### Generating barcoded destination vector

Barcoded plasmid constructs (pMV2_BC, Addgene_262837) were prepared by cloning barcode sequences into the pMV2 backbone (pMV2, Addgene_263394), followed by transformation into Electro-Competent ccdB survival T1R cells as described in the protocol available on protocols.io (doi: 10.17504/protocols.io.m7vznm1xrvx1/v1). The protocol for making the Electro-Competent ccdB survival T1R cells is available in (Aslanzadeh et al., 2026), and on protocols.io (doi: 10.17504/protocols.io.qpvrex7bzlmk/v1). We estimate approximately 20 million unique barcodes in the destination vector based on colony counts from dilution plates and sequencing of individual colonies.

#### Oligonucleotide pool design and preparation

Degenerate oligonucleotides combining NNS and NNK codons were designed using the SUNi primer design method (Mighell et al., 2023), implemented in MOLigos_Script_GBA1.ipynb (available on GitHub at https://github.com/vaslanzadeh/gba1_flowseq_pub and archived on Zenodo, https://doi.org/10.5281/zenodo.24116674). Oligonucleotide design was performed as described in the protocol available on protocols.io (doi: 10.17504/protocols.io.92ld3b6bxg5b/v1). This allowed a single amino acid mutation to be introduced at each of the 536 codons across the *GBA1* coding sequence (excluding the start codon) (Supplementary Table 6). Together, the NNS/NNK codons encode all 20 amino acids plus one stop codon. To avoid uneven representation of variants, oligonucleotides were split across 10 pools, each spanning 55 consecutive codons, with mutagenesis carried out separately for each pool. The mutagenesis primers and the universal secondary primer (VA03) (Supplementary Table 6) were phosphorylated in preparation for saturation mutagenesis as described in the protocol available on protocols.io (doi: 10.17504/protocols.io.6wgq9re6olk5/v1).

#### Variant library generation by saturation mutagenesis

Nicking mutagenesis protocol based on (Wrenbeck et al., 2016) followed to generate the *GBA1* variant library and was performed as described in the protocol available on protocols.io (doi: 10.17504/protocols.io.6wgq9re6olk5/v1). Mutagenesis reactions were performed independently for each of the 10 pools using a 1:500 dilution of the corresponding phosphorylated oligos. Following mutagenesis, the reaction products were purified, transformed into high-efficiency NEB 5-alpha competent *E. coli*. Based on the serial dilutions, transformation efficiency was estimated at approximately 4.97 × 10⁵ CFU per pool. The purified plasmid was used as the entry plasmid for LR Gateway cloning to generate the barcoded *GBA1* variant library.

#### Generating barcoded *GBA1* variant library by LR Gateway cloning

The pAINT-*GBA1* variant library was cloned into the barcoded pMV2 plasmid (Addgene_262837) with Gateway LR recombination as described in the protocol available on protocols.io (doi: 10.17504/protocols.io.8nlk5b625l5r/v1). Separate LR reactions were performed for each of the 10 pools. For each reaction, 150 ng of entry plasmid (*GBA1* variant library) and 150 ng of destination plasmid (barcoded pMV2 plasmid) were assembled with Invitrogen Gateway LR Clonase II enzyme mix (Thermo Fisher Scientific, 11791020). For each pool, two independent transformations were carried out for 10uL of LR product. To assess transformation efficiency and colony number, a 50 µL aliquot of each transformation mixture was taken for plating onto a 100 µg/mL ampicillin LB-agar plate. Based on the colony counts, each pool yielded approximately 10,000–22,000 CFU. Equal proportions of purified plasmid from each pool were combined to make the final *GBA1* variant barcoded library (pMV2-GBA1, Addgene_263443).

#### PacBio sequencing of barcoded *GBA1* variant library

To link each barcode to its corresponding *GBA1* variant, we carried out PacBio long-read sequencing. Samples were prepared for sequencing as described in the protocol available on protocols.io (doi: 10.17504/protocols.io.p2l64kpbrvqe/v1). Sequencing templates were prepared from the purified plasmid library through triple digestion with AscI, AgeI-HF, and BssSI-v2, which yielded a 2091 bp fragment containing the barcode and *GBA1* sequence, along with 600-700 bp and 1.3-1.4 kb fragments derived from the plasmid backbone. Following purification of the digestion products, samples were submitted to Edinburgh Genomics (University of Edinburgh) for size selection on a BluePippin (Sage Science), then library preparation and sequencing on a single SMRT Cell using the PacBio Revio platform. HiFi reads (GEO; GSE346421) were processed with alignparse (Crawford & Bloom, 2019) version 0.6.3, to align reads and call mutations within the *GBA1* sequence and link these mutations with their barcodes. Of 8,664,862 HiFi reads, 7,410,187 mapped to the target and used to build a codon-variant lookup table.

Reads were first filtered to retain only mapped HiFi reads with a PacBio sequencing accuracy of at least Q40 across both the *GBA1* sequence and the barcode. Because PacBio sequencing yielded an average of 23.16 reads per amplicon (Supplementary Table 7), we then determined the empirical accuracy of the retained reads (Starr et al., 2020) to evaluate how reliably barcodes were phased with variants, in other words, how consistently reads sharing a given barcode agreed with one another. Excluding reads containing indels, empirical accuracy per HiFi read was 0.99. Given that 94.1% of barcodes (191833) had two or more associated HiFi reads, a cumulative consensus was built from the HiFi reads within each such barcode, raising consensus accuracy above the empirical accuracy calculated for single reads. Barcodes linked to insertions, deletions, or more than one variant were excluded from further analysis. In total, 184878 barcodes were retained: 123668 corresponded to missense variants, 4896 to nonsense variants, and 7182 and 54028 to synonymous variants and wild-type sequence, respectively (Supplementary Table 7). The code used to build the final barcode-variant table is available on GitHub (https://github.com/vaslanzadeh/gba1_flowseq_pub) and archived on Zenodo (https://doi.org/10.5281/zenodo.24116674).

#### Library transfection

The transfection of the *GBA1* variant barcoded plasmid library into *GBA1*-fKO HEK293 Bxb1 landing pad cells was performed as described in the protocol available on protocols.io (doi: 10.17504/protocols.io.p2l64km8zvqe/v1). Following transfection, puromycin selection was applied to select for stable, single-copy integrants. Cells were maintained under puromycin selection, expanded sequentially in T175 and T25 flasks for two weeks before harvesting.

#### Genomic DNA extraction, barcode amplification and sequencing

Singlets were sorted into four bins according to fluorescence-intensity using BD FACS Aria II, collecting a minimum of 10 million cells per bin. The sorted cells were split into four 1.5 mL aliquots per bin, pelleted, and stored at −20 °C. Genomic DNA extracted with phenol:chloroform:isoamyl alcohol and subsequent barcode amplification was performed as described in the protocol available on protocols.io (doi: 10.17504/protocols.io.m7vznm16rvx1/v1). Following precipitation, the four tubes per bin were combined to yield 400 μL of purified genomic DNA per population. Genomic DNA barcodes were amplified via PCR using an indexed Illumina P7 forward primer (VA05) and an Illumina P5 reverse primer (VA04), which is complementary to a genomic region downstream of the integration site, to ensure specific amplification and prevent the unwanted amplification from residual pMV2-*GBA1* plasmid DNA. The purified product was sent to WTCRF for size selection of the 288 bp product, followed by equimolar mixing. Sequencing was performed on Illumina NextSeq sequencing with a custom primer (VA06). Demultiplexed sequencing reads were trimmed to the 30-nucleotide barcode region using Trimmomatic. Enrich2 was then used to align the trimmed barcodes against the defined barcode library based on the PacBio sequencing result and generate counts across all FACS bins. Raw and processed sequencing data are available on GEO (GSE346421), and analysis scripts are available on GitHub (https://github.com/vaslanzadeh/gba1_flowseq_pub) and archived on Zenodo (https://doi.org/10.5281/zenodo.24116674).

#### Calculation of variant function scores and empirical p-values

Using the barcode counts obtained above, variant function scores and empirical p-values were calculated using a previously published method (Aslanzadeh et al., 2025), with a small modification described below. Briefly, barcode frequencies within each FACS bin were calculated from Enrich2 counts, and a weighted barcode score was derived from each barcode’s frequency distribution across the four fluorescence bins. Barcode scores were averaged per variant across barcodes and replicate experiments to yield a mean log2 variant score. To assess whether a variant’s score differed significantly from wild-type, we performed a bootstrapping procedure in which weighted averages of randomly sampled sets of wild-type barcodes, matched in number to the barcodes available for each variant, were used to generate an empirical null distribution, from which an empirical p-value was calculated for each variant.

We note that in our previous publication (Aslanzadeh et al., 2025), these empirical p-values were referred to as false discovery rates (FDR); because no multiple-testing correction was applied, this terminology was inaccurate, and we refer to them as empirical p-values throughout the present manuscript.

#### Generating single substitution-stable cell lines

Single-variant stable cell lines expressing WT *GBA1* or 24 individual *GBA1* mutants were generated in stable cell lines (Table 1). Mutant primers were designed for each mutant and ordered from IDT (Supplementary Table 6). *GBA1* mutant constructs were generated via site-directed mutagenesis, and each variant was transfected into a *GBA1* fKO HEK293 Bxb1 landing pad cell line, as described in the protocol available on protocols.io (doi: 10.17504/protocols.io.4egnxbn4ql5d/v1). Individual plasmids were confirmed by sequencing with Plasmidsaurus. Flow cytometry was performed on the stable cells to ensure BFP knockout, indicating effective stable integration.

### Analysis of clinically annotated variants

#### MDSGene Database Annotated Variants

*GBA1* variants were catalogued, with genetic annotations assigned by MDSGene according to ACMG criteria and clinical severity annotations obtained from MDSGene using data from the GBA1 PD Browser, as previously described (Rossi et al., 2025). Missense and nonsense variants with corresponding flow-seq scores were included in the analysis.Variants supported by at least 3 barcodes and with an *p* ≤ 0.05 were classified as functionally deleterious. Variants supported by fewer than three independent barcodes were considered to have insufficient confidence for functional interpretation and were indicated with unfilled circles. Plots were generated in R using the ggplot and ggbeeswarm packages.

#### Global Parkinson’s Genetics Program (GP2)

We used whole-genome sequencing (WGS) data from GP2 Release 12 (https://doi.org/10.5281/zenodo.20932193). *GBA1* variants were extracted from PLINK files using PLINK and annotated using ANNOVAR (K. Wang et al., 2010); in addition, the Gauchian pipeline (Toffoli et al., 2022) was used for *GBA1*-specific variant calling. In contrast to the complete set of theoretically possible variants evaluated in the MAVE assay, analyses were restricted to *GBA1* variants that are possible based on single-nucleotide substitutions in the human coding sequence.

Genetic association and age at onset (AAO) analyses were restricted to the European ancestry dataset, which provided sufficient sample size for meaningful variant-level analyses. Single-variant association analyses were performed using PLINK, including sex and the first five genetic principal components as covariates. Odds ratios (ORs) were considered for correlation with MAVE scores only for variants observed in >5 or >10 individuals across PD cases and controls, with at least one carrier required in each group. AAO values were obtained from GP2 clinical data, and analyses were restricted to individuals carrying a single heterozygous *GBA1* variant; individuals with homozygous variants or two different heterozygous *GBA1* variants were excluded. Associations between MAVE scores and genetic effect estimates or AAO were assessed using Spearman’s rank correlation coefficient, and plots were generated using Matplotlib (Hunter, 2007) and Seaborn (Waskom, 2021) in Python.

#### Comparison with computational variant-effect predictors and evolutionary conservation

To evaluate the concordance between the experimental variant scores and established pathogenicity predictors (Supplementary Table 4), correlation analyses were performed using Python (via the pandas and scipy libraries). Spearman rank correlation was calculated for each pair of variables, excluding variants with missing data for a given predictor.

For visualization, scatter plots were generated using the matplotlib and seaborn packages. A linear regression line of best fit was overlaid on each plot to illustrate the general trend. To provide clinical context, all plotted variants were color-coded according to their pathogenicity classifications based on MDSGene. Finally, a predefined subset of 20 clinically relevant variants (including N409S and L483P) was annotated on the plots.

To assess the relationship between Flow-seq scores and evolutionary conservation, the median Flow-seq score was calculated for each residue position using variants supported by ≥3 barcodes. Evolutionary conservation scores for the amino acid residues were calculated using the ConSurf web server (https://consurf.tau.ac.il/) (Yariv et al., 2023). These position-level median Flow-seq scores were then compared with the corresponding ConSurf conservation scores. Plots were generated in R using the ggplot package.

## Data and Code Availability

The data, code, protocols and key lab materials used and generated in this study are listed in a Key Resources Table (KRT) with their persistent identifiers, available at Zenodo (https://doi.org/10.5281/zenodo.23169564). Raw and processed experimental data, including barcode and variant scores, Enrich2 count tables, raw flow cytometry (FCS) files, immunoblot images and quantification, and a README describing every file, are deposited at Zenodo (https://doi.org/10.5281/zenodo.23169564). Raw sequencing data are available in NCBI GEO under accession GSE346421. The variant scores are available in MaveDB (urn:mavedb:00001327-a; https://www.mavedb.org/experiments/urn:mavedb:00001327-a) and can be explored interactively at https://gba1-app.onrender.com/. All code used to process and analyse the MAVE data and to generate the figures is available on GitHub (https://github.com/vaslanzadeh/gba1_flowseq_pub) and archived on Zenodo (https://doi.org/10.5281/zenodo.24116674). Protocols are available on protocols.io (see Methods and KRT for individual DOIs).

Data used in the preparation of Figure 4 were obtained from the Global Parkinson’s Genetics Program (GP2; https://gp2.org). Specifically, we used Tier 2 data from GP2 release 12 (https://doi.org/10.5281/zenodo.20932193). GP2 data can be requested through AMP PD (https://amp-pd.org). All code generated for the GP2 analyses is available on GitHub (https://github.com/GP2code/GBA1-MAVE) and archived on Zenodo (https://doi.org/10.5281/zenodo.23654228).

## Supplementary Material

**Supplementary Figure 1:**
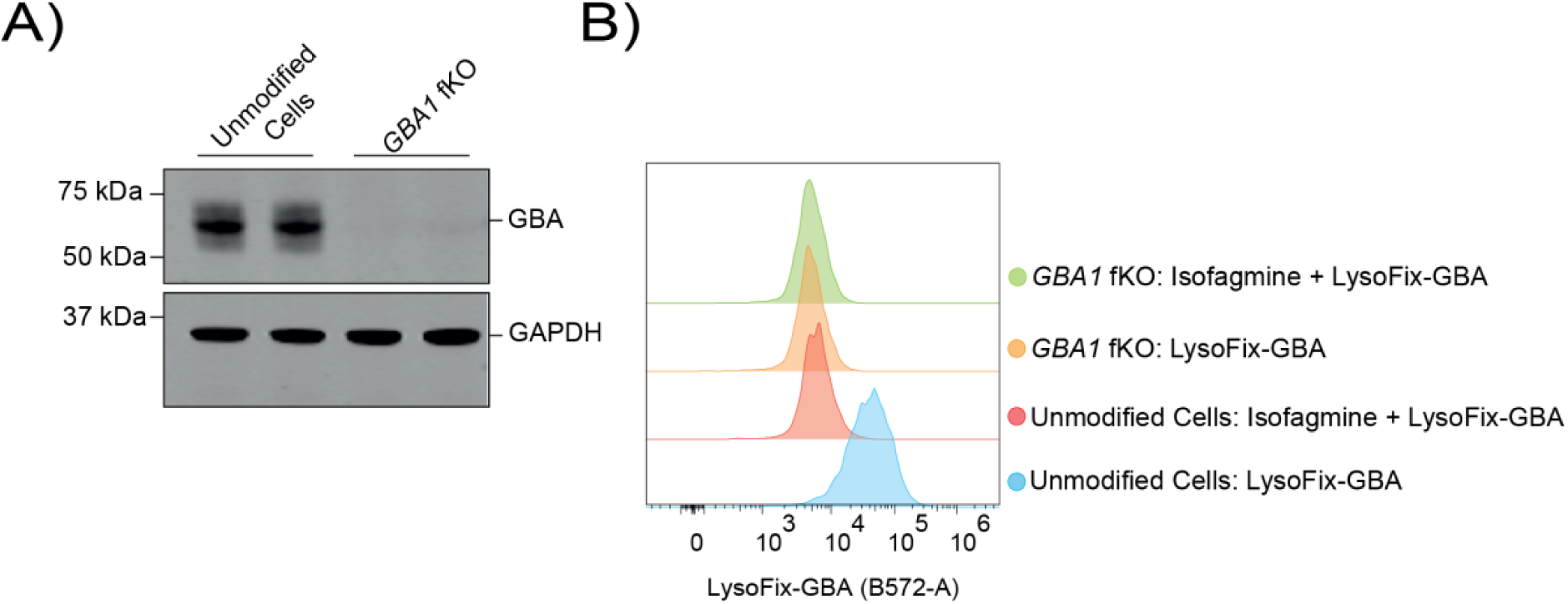
Validation of *GBA1* total protein expression and activity in a *GBA1 f*KO cell line. A) Representative western blot of total *GBA1* in an unmodified HEK Bxb1 landing pad line and a *GBA1* fKO HEK Bxb1 landing pad line. B) Representative flow cytometry histogram plots for LysoFix-GBA treated *GBA1* fKO or unmodified HEK Bxb1 landing pad cells. Cells were pre-treated with 10 µM Isofagomine or vehicle control (DMSO) for 1 hour, followed by treatment with 15 µM LysoFix-GBA for 1 hour.

**Supplementary Figure 2:**
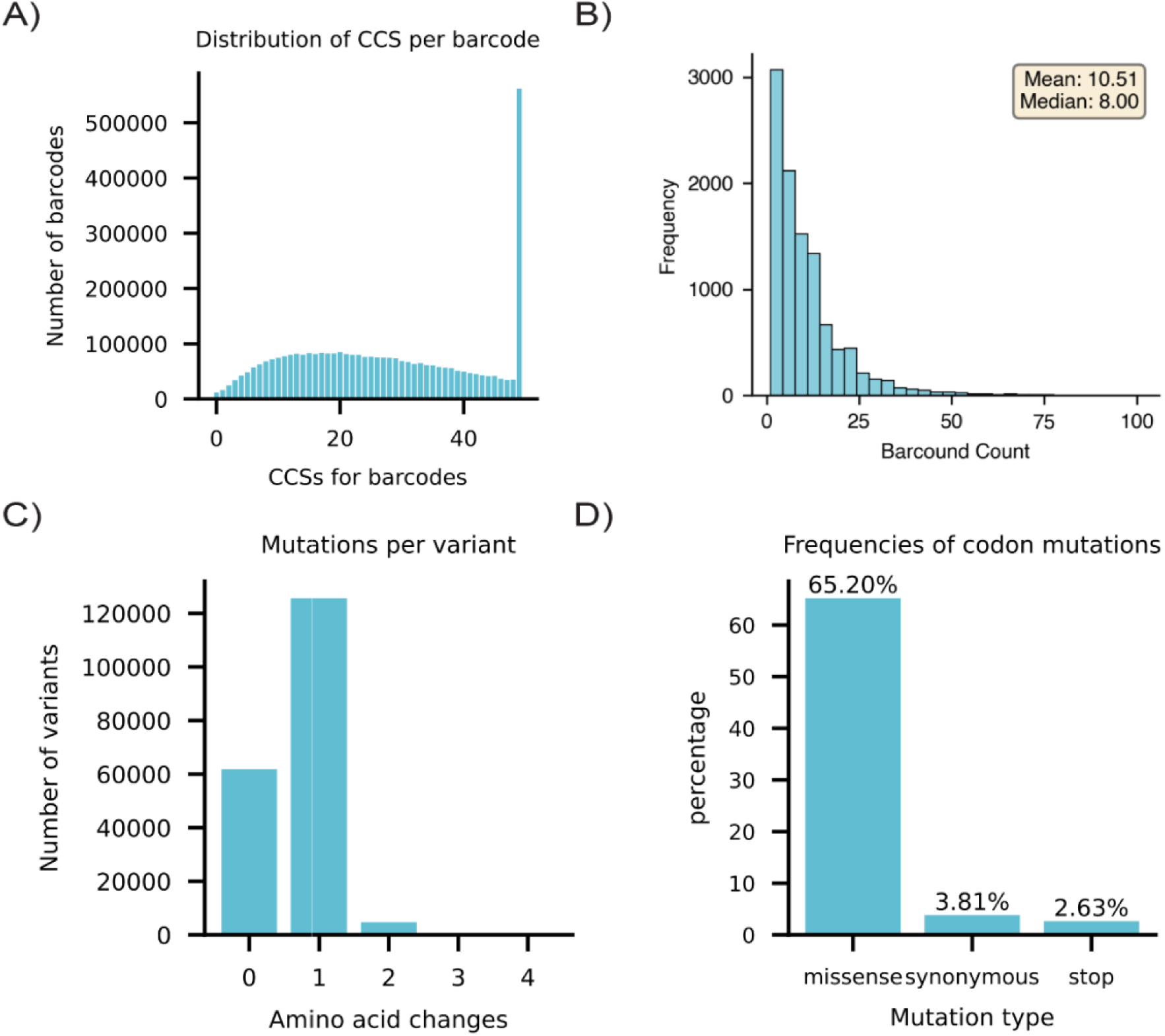
Composition of *GBA1* variant construct library and barcode scores. A) Number of circular consensus sequences (CCSs) generated by PacBio sequencing and used for barcode-variant phasing. B) Distribution of barcode counts per *GBA1* variant. C) The number of amino acid changes per *GBA1* variant. D) The percentage of missense, nonsense, and synonymous *GBA1* variants across the plasmid library.

**Supplementary Figure 3:**
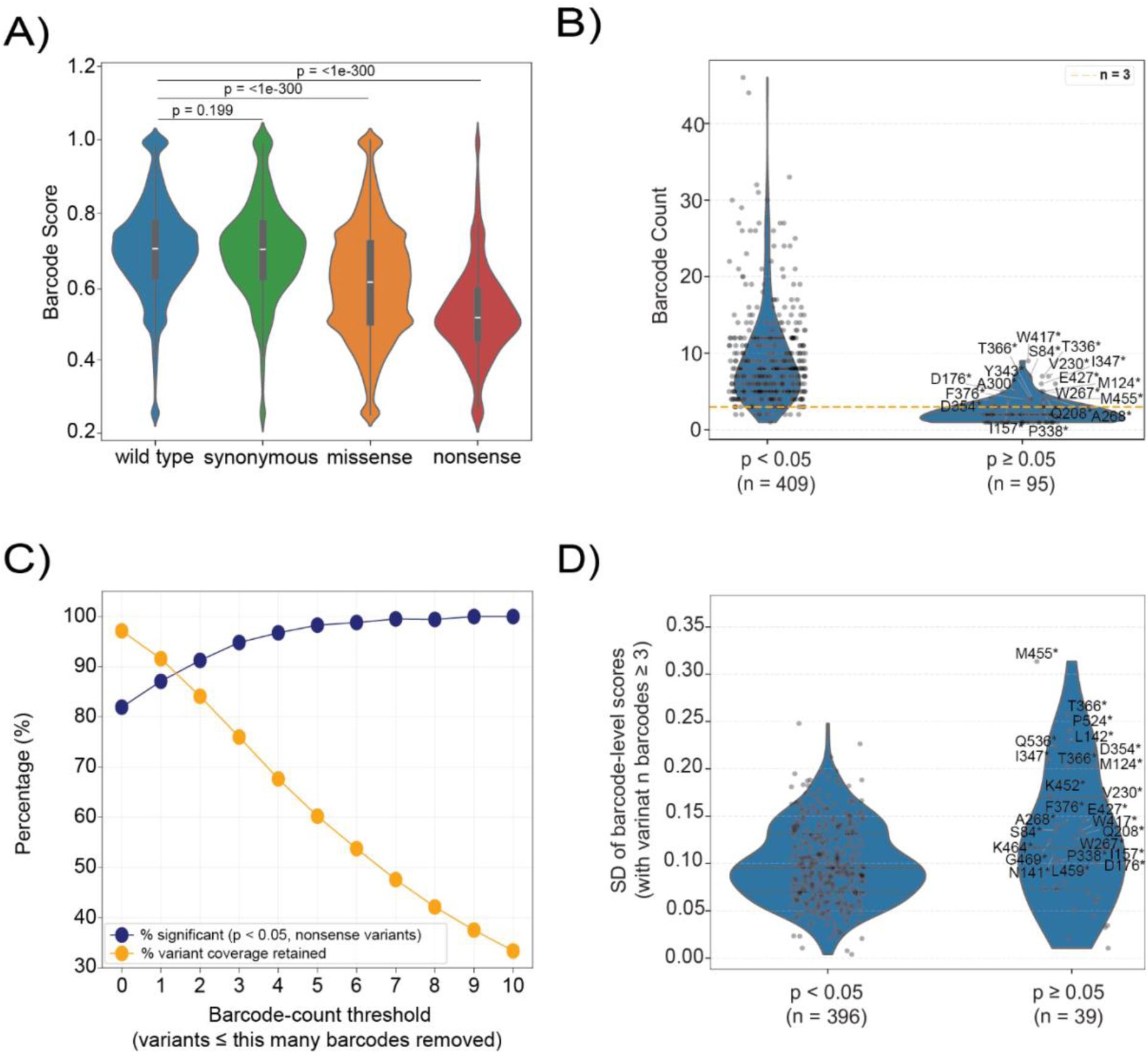
Distribution of flow-seq scores and barcode coverage across variant classes. **A)** Violin plot showing the distribution of flow-seq barcode scores stratified by variant consequence class; wild-type (n = 47,733), synonymous (n = 6326), missense (n = 105,245), and nonsense (n = 4338). Statistical significance was assessed using two-sided Mann-Whitney U test, with the overall comparison between wild-type and missense or nonsense yielding *p* < 1 × 10⁻³⁰⁰. B) Barcode counts for nonsense variants (downstream of residue 21) stratified by empirical p-value significance, *p* < 0.05 (n = 409) and *p* ≥ 0.05 (n = 95). A barcode count of 3 is indicated by a yellow line. Barcode counts were compared using a two-sided Mann–Whitney U test (*p* = 2.37 × 10⁻^39^). C) The percentage of nonsense variants (downstream of residue 21) retained as a function of the barcode coverage threshold. At each threshold, variants with ≤ the specified number of barcodes were excluded from the analysis. The percentage of retained variants is indicated in blue, and the percentage of retained variants with a score significantly different from WT (*p* < 0.05) is indicated in orange. D) Distribution of the standard deviation (SD) of barcode flow-seq scores for nonsense variants with barcode coverage ≥3, stratified by empirical p-value significance, *p* < 0.05 (n = 395) and *p* ≥ 0.05 (n = 40).

**Supplementary Figure 4:**
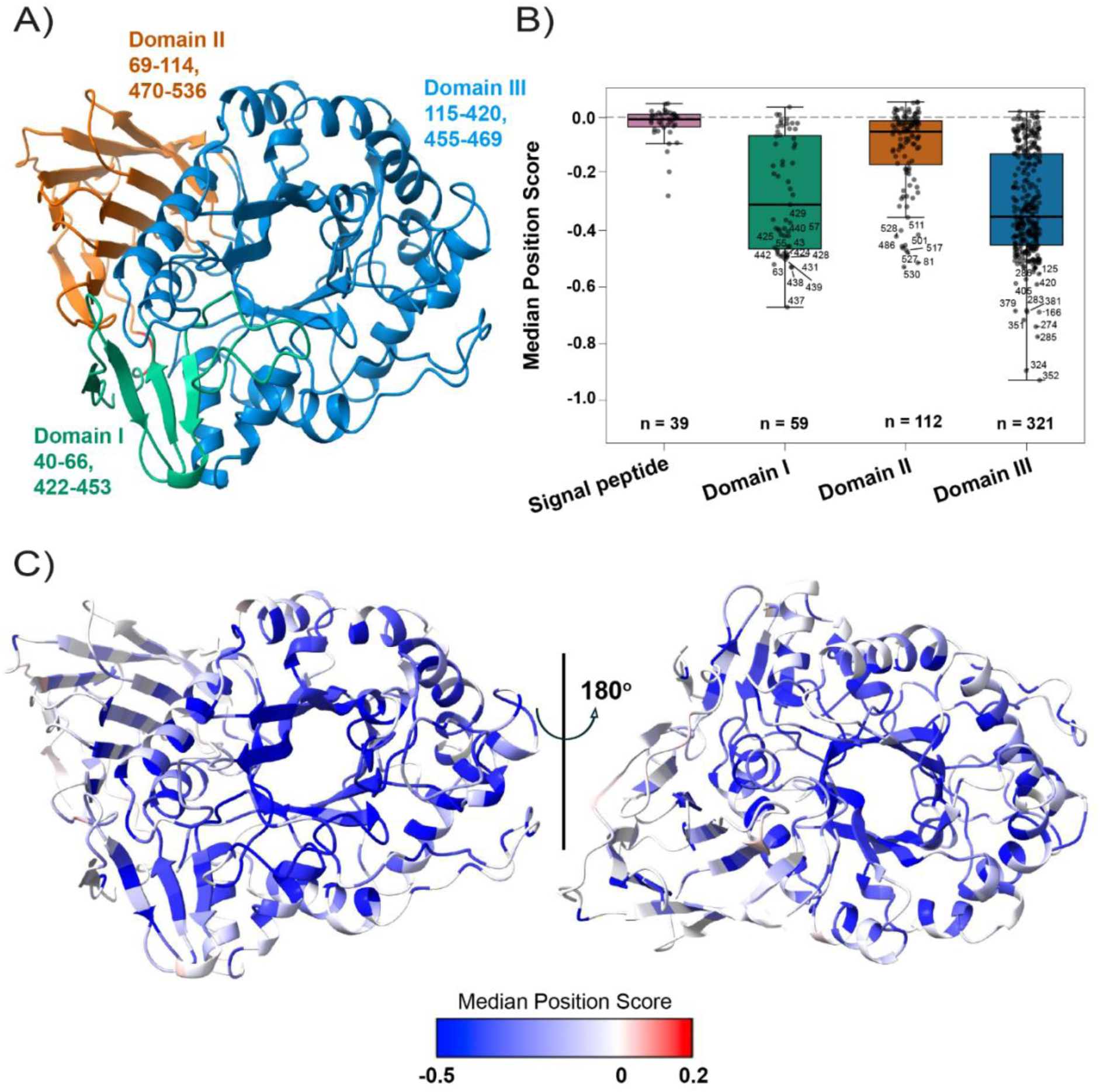
Structural mapping of median flow-seq scores onto *GBA1*. A) the mature *GBA1* structure following cleavage of the 39-amino-acid N-terminal signal peptide (PDB 2NSX). The mature protein comprises three structural domains: domain I (residues 40–66 and 422–453, green), domain II (residues 69–114 and 470–536, orange), and domain III (residues 115–420 and 455–469, blue) (Dvir et al., 2003; Kacher et al., 2008). B) Median position flow-seq score across domains. Each point represents the calculated median flow-seq score of variants at each position. C) *GBA1* structure shown in two orientations, with each residue coloured according to its median flow-seq score. Scores are shown relative to WT *GBA1*, with WT-like median scores shown in white, gain-of-function median scores in red, and loss-of-function median scores in blue.

**Supplementary Figure 5.**
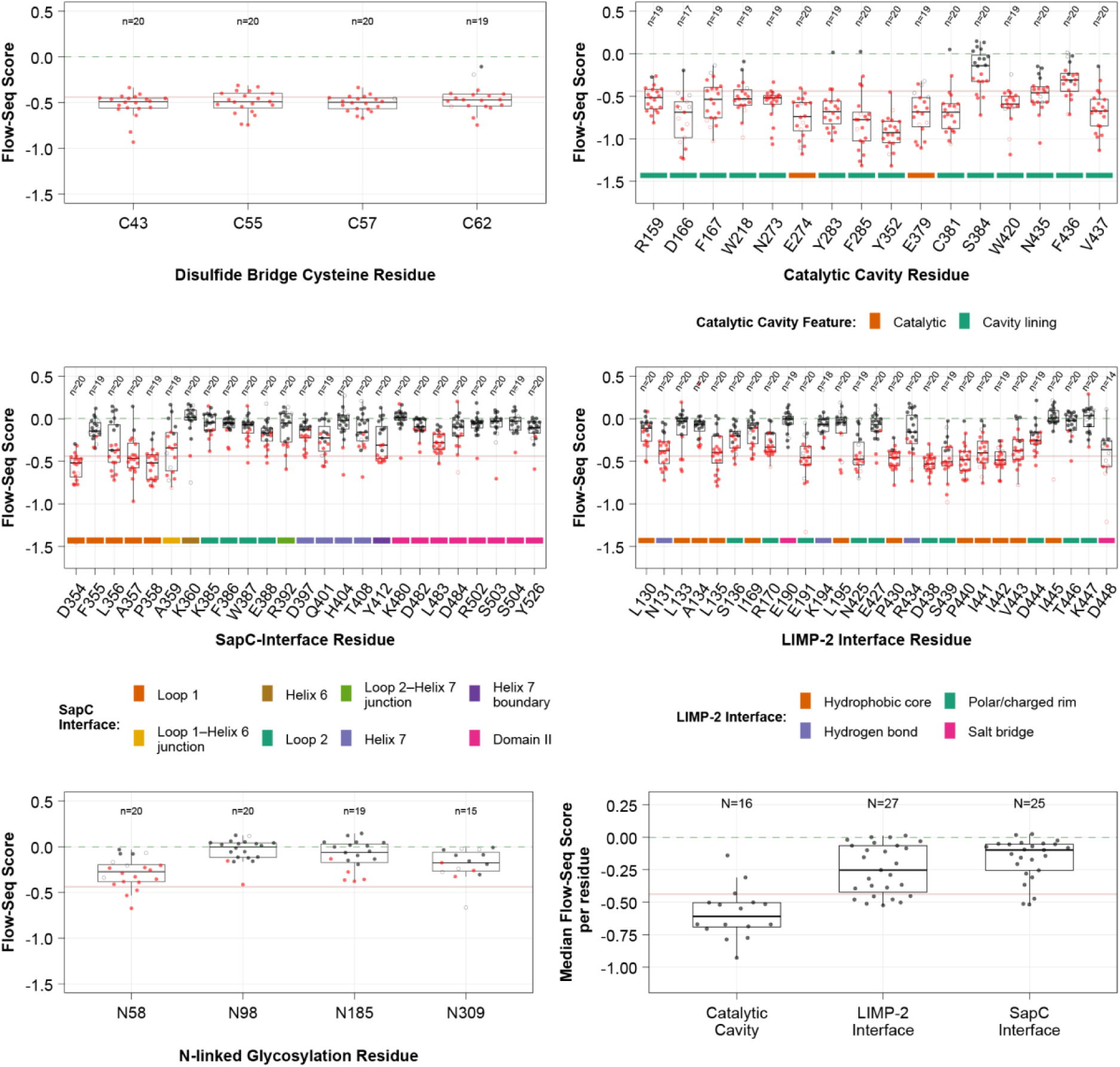
Flow-Seq scores across *GBA1* structural regions. Distribution of Flow-Seq scores for *GBA1* variants located within proposed structural regions: (A) domain I disulfide bridge (Dvir et al., 2003), (B) catalytic cavity (Lieberman, 2011), (C) SapC binding interface (Romero et al., 2019), (D) LIMP-2 binding interface (Dobert et al., 2025; Liou et al., 2014), and (E) N-linked glycosylation sites (Berg-Fussman et al., 1993). Each point represents a variant at the indicated residue position. Red points indicate variants with a *p* < 0.05, while filled and unfilled points represent variants supported by ≥ 3 barcodes or < 3 barcodes, respectively. F) Median flow-seq score of residues associated with the catalytic cavity, LIMP-2 interface or SapC interface. Each point represents the calculated median flow-seq score of variants at a given position. Residues assigned to each structural group are provided in Supplementary Table 3.

**Supplementary Figure 6.**
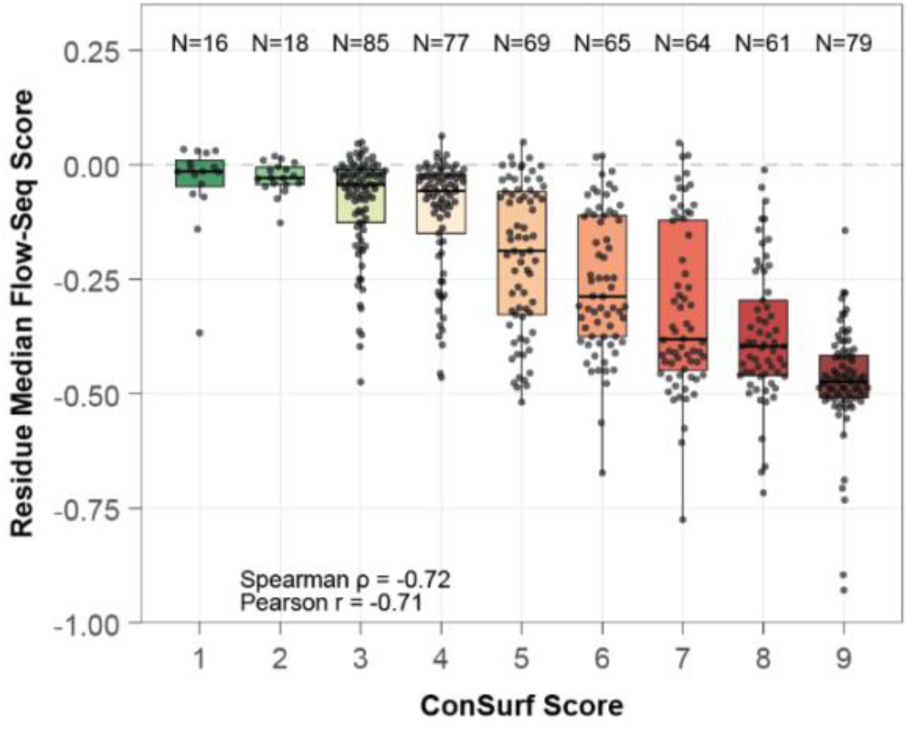
The relationship between evolutionary conservation and variant functional effects. Distribution of flow-Seq scores grouped by ConSurf conservation score. Each point represents the calculated median flow-seq score of variants at each position.

**Supplementary Figure 7:**
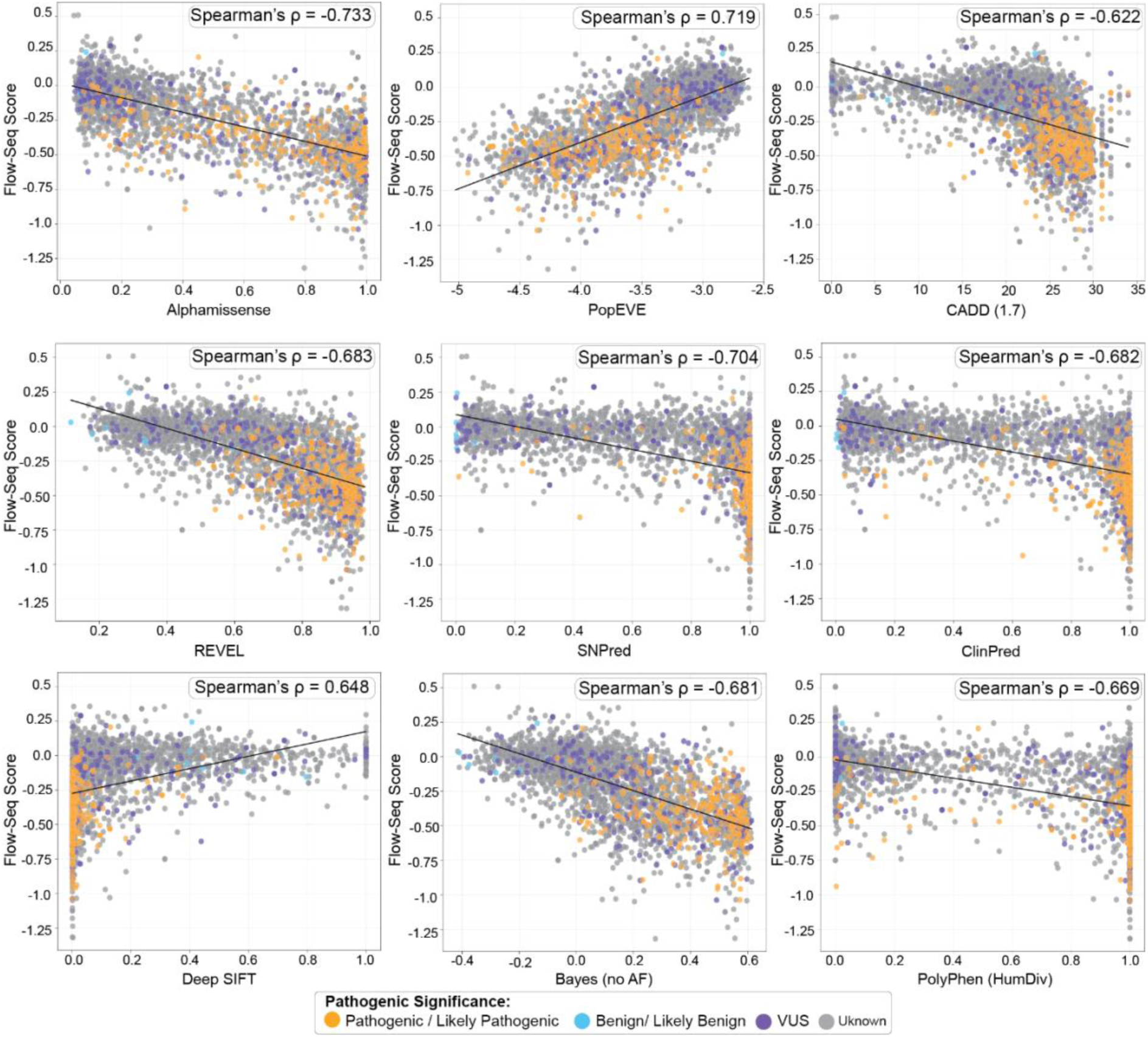
Comparison of experimental functional flow-seq scores with computational predictions of variant effect. Experimental functional scores are plotted against scores from the VEPs, AlphaMissense, PopEVE, CADD (v1.7), REVEL, SNPred, ClinPred, DeepSIFT, Bayes no Allele Frequency (no AF), and PolyPhen-2 (HumDiv). Spearman’s rank correlation coefficients were calculated to assess the relationship between each computational prediction score and the functional score. Variants are coloured according to MDSGene genetic annotation: pathogenic/likely pathogenic (orange), benign/likely benign (blue), variants of uncertain significance (VUS; purple), and unknown (grey).

**Supplementary Figure 8:**
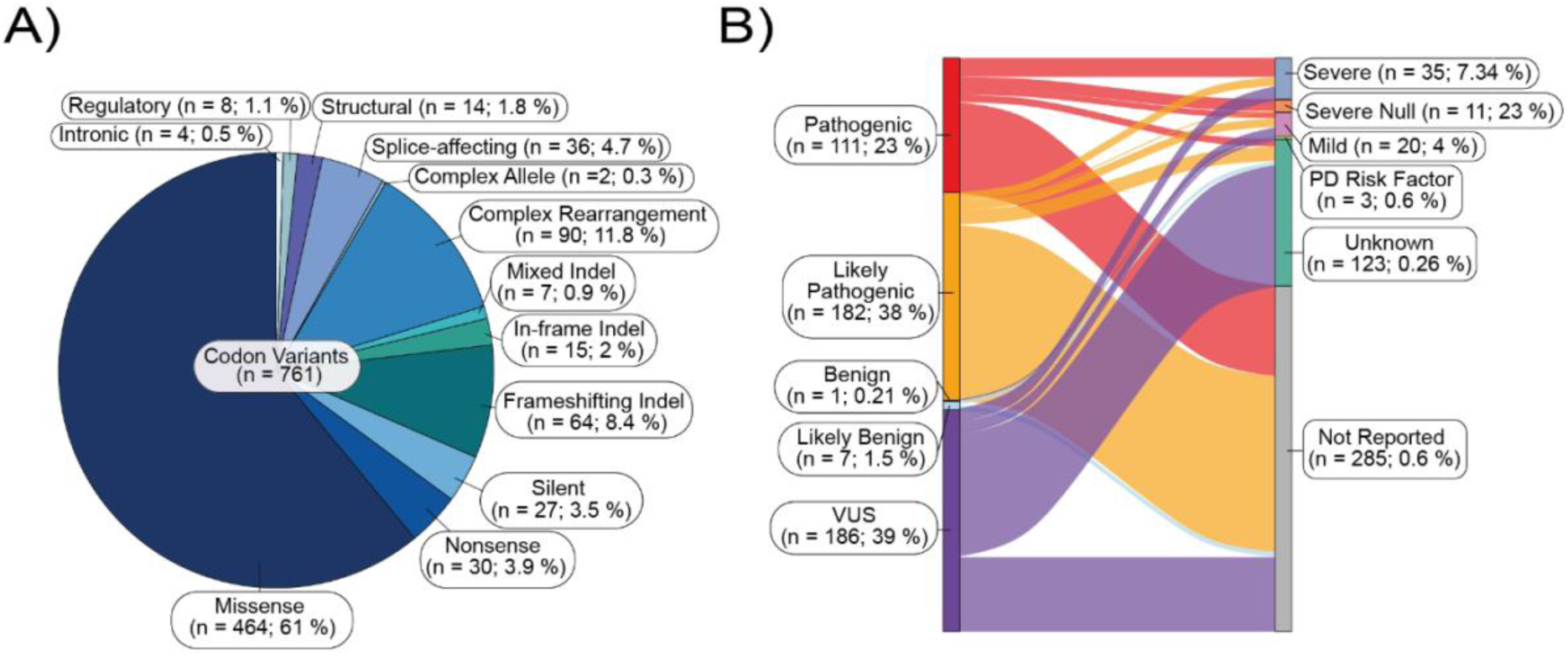
Distribution of *GBA1* variants reported in the MDSGene database. (A) Pie chart showing the distribution of unique *GBA1* codon variant consequence classes among *GBA1* variants reported in the MDSGene database. A total of 761 unique codon-level variants were identified, including 464 missense and 30 nonsense variants. (B) Sankey plot illustrating the relationship between pathogenicity classification and clinical severity among the 477 protein-coding *GBA1* variants reported in the MDSGene database for which Flow-seq functional scores were available.

**Supplementary Figure 9.**
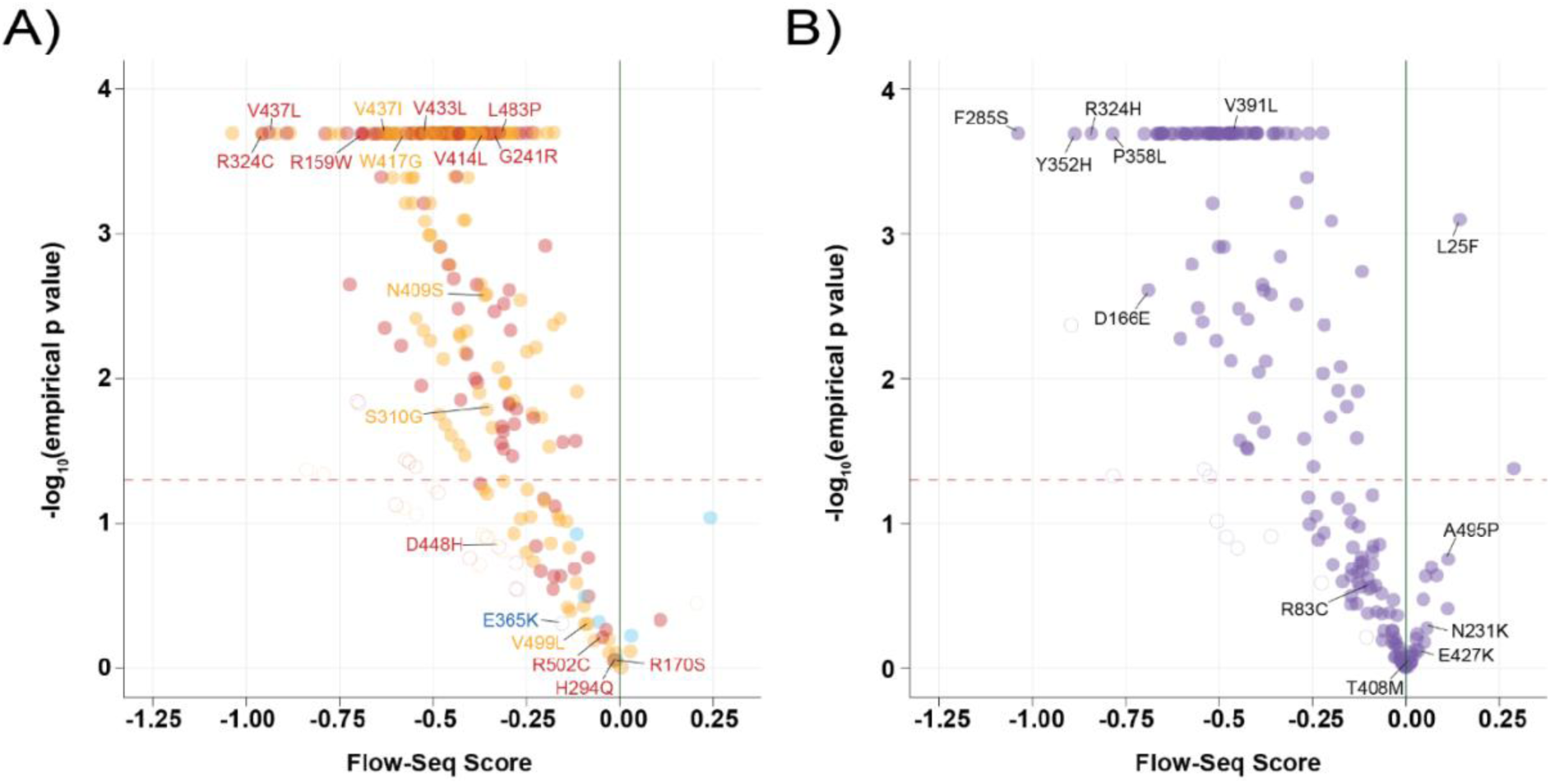
Functional effects of MDSGene-annotated variants. Volcano plots showing flow-seq variant score versus −log10(*empirical p-value*) for 477 protein coding *GBA1* variants annotated in the MDSGene database. (A) Variants annotated as pathogenic (red, n = 111), likely pathogenic (orange, n = 172), benign (dark blue, n = 1), or likely benign (light blue, n = 7). (B) Variants annotated as VUS (purple, n = 186). The horizontal dashed red line indicates the significance threshold (p < 0.05), and the solid green vertical line indicates the WT reference score (flow-seq score = 0). Filled and open circles denote variants with ≥ 3 barcodes or < 3 barcodes respectively.

**Supplementary Figure. 10:**
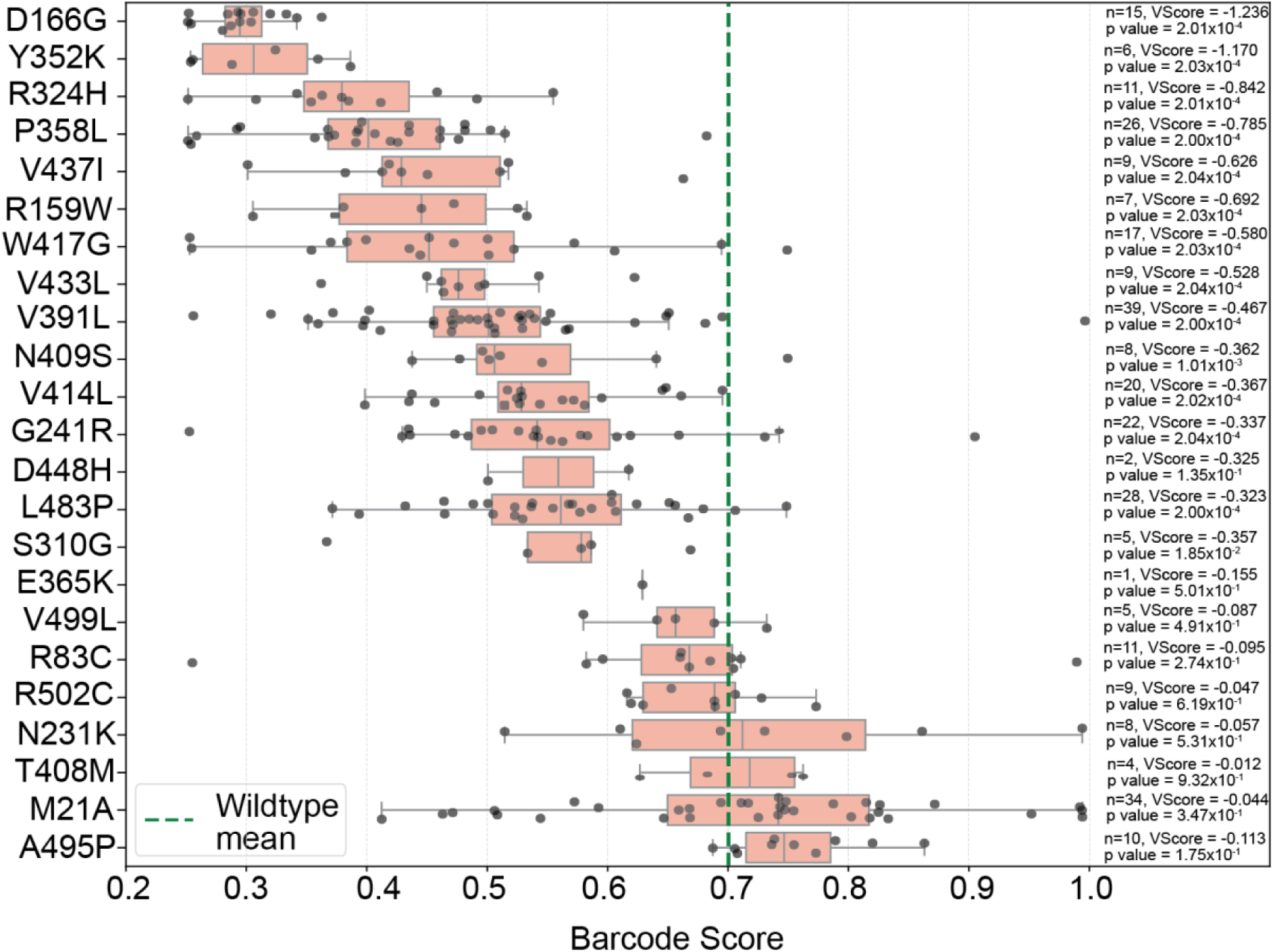
Distribution of barcode scores for selected variants. Individual data points correspond to variant scores (VScore) derived from unique barcodes. Green dashed line indicates average WT barcode score for.

**Supplementary Figure. 11:**
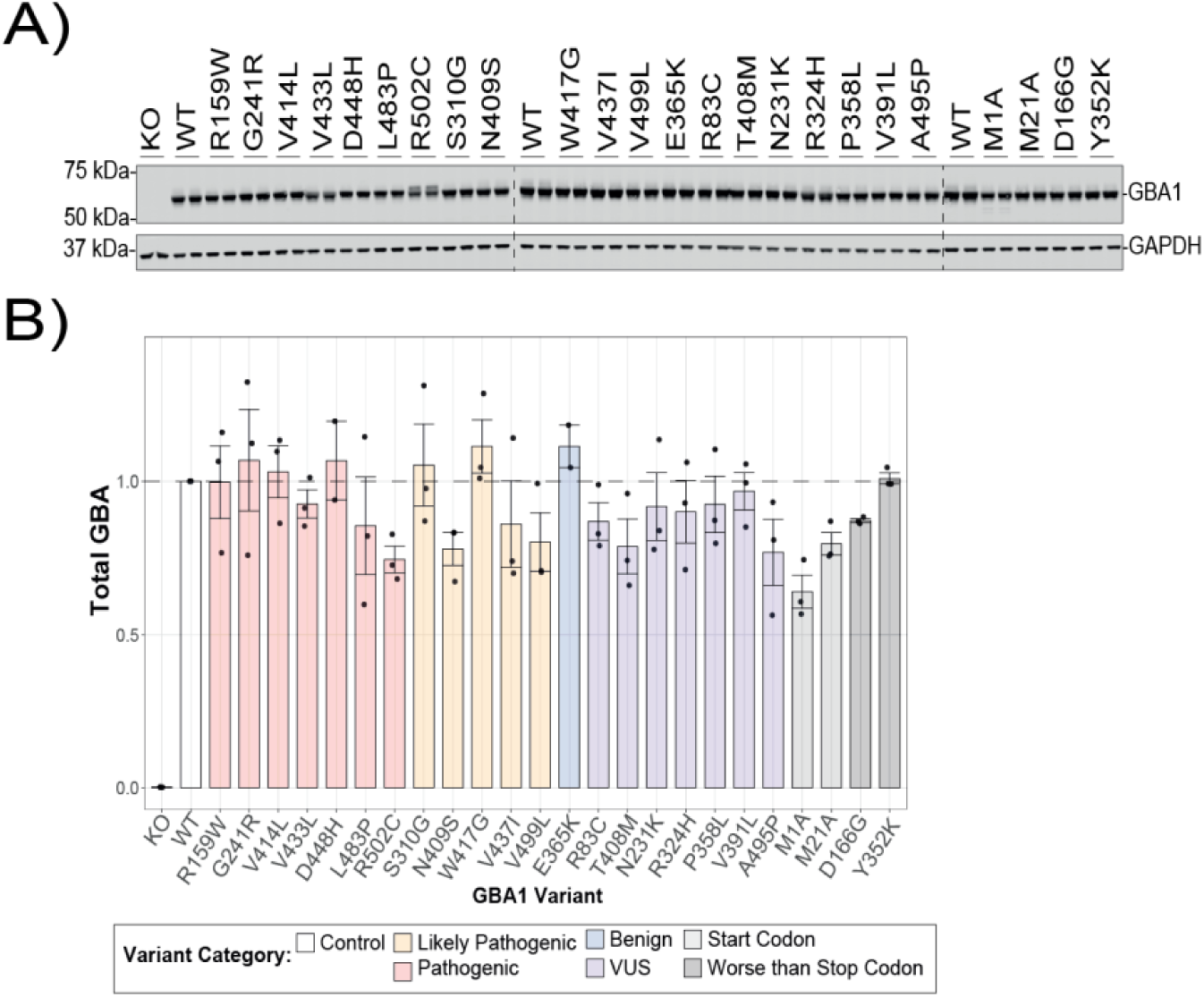
Quantification of total *GBA1* protein levels in single-variant HEK Bxb1 landing pad cell lines. (A) Representative immunoblot showing total *GBA1* protein levels in single-variant expressing cell lines. (B) Quantification of total *GBA1* protein levels from three independent experiments. No significant differences were observed between variants and control. Data were analyzed using one-way ANOVA followed by Dunnett’s multiple comparisons test. Error bars represent mean ± SEM. Variants were coloured according to category: pathogenic (red), likely pathogenic (orange), benign (dark blue), and VUS (purple), start codon (light grey), worse-than-stop-codon (dark grey), and control (white).

**Supplementary Figure 12.**
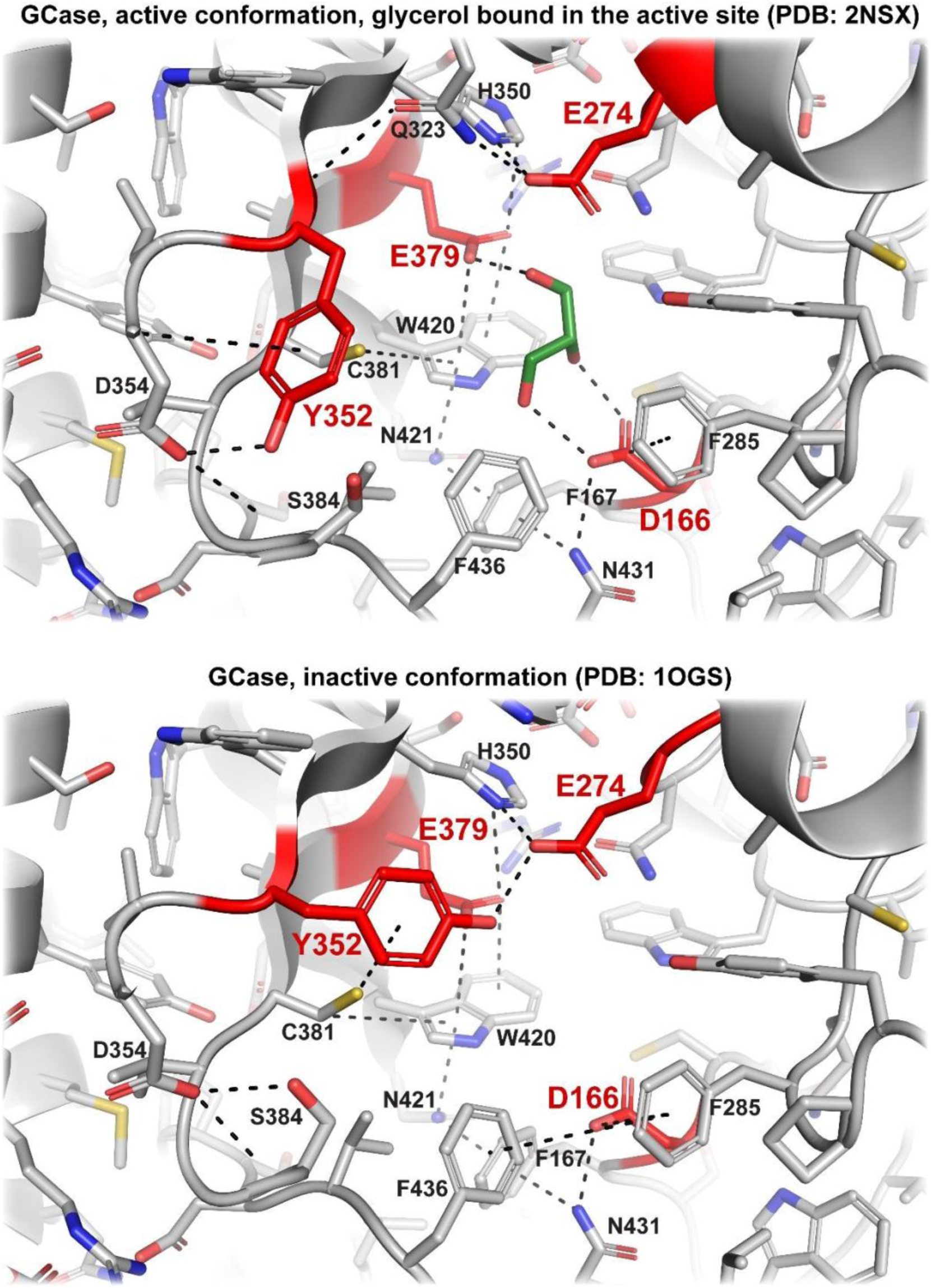
Structural modeling of the GCase catalytic cavity. Structural mapping of GCase in the active conformation (PDB: 2NSX) and in the inactive conformation (PDB: 1OGS). In the active conformation, glycerol is shown in green within the catalytic site, highlighting interactions within the substrate-binding region. Residues E274 and E379 (shown in red) engage the substrate through complementary hydrogen-bonding and electrostatic interactions, with E274 functioning as the general acid/base and E379 serving as the catalytic nucleophile. Residue Y352 (red) contributes to the loop rearrangement between the extended (inactive) and helical (active) conformations, while D166 (red) contributes to substrate binding.

**Supplementary table 1:** *GBA1* construct library score summary.

|  |  |
|---|---|
| number of ccs reads (HiFi Reads) | 8664862 |
| median ccs read quality | Q40.0 |
| ccs aligned to the amplicon | 7410187 |
| low qualtiy or unmapped ccs (filtered) | 1254675 |
| average (median) ccs per barcode | 23.16 |
| total number of barcodes with consensus sequences | 192377 |
| barcodes with consensus seqeunces (no_indels with zero or single aa changes) | 184878 |
| barcodes with single_aa_substitutions (nonsynonymous) | 123668 |
| barcodes with single_aa_substitutions (stop codon) | 4896 |
| barcodes with synonymous mutations | 7182 |
| barcodes with WT sequence, no mutagenesis | 54028 |
| unique aa substitutions in the library (library completeness) | 10460 |
| missense aa substitutions associated with single barcode | 500 |
| missing missense single aa substitutions from the library (zero barcode) | 286 |
| unique stop variants in the library | 526 |

**Supplementary table 3:** Summary of Structural Annotation References.

| Site | GBA1 Feature | Residue | Residue Alias | Reference |
|---|---|---|---|---|
| LIMP-2 Binding Interface | Hydrophobic core interface | L130, L133, A134, L135, I169, L195, P430, P440, I441, I442, V443, I445 | L91, L94, A95, L96, I130, L156, P391, P401, I402, I403, V404, I406 | Dobert et al., 2025 (PBD; 9FJF). Liou et al., 2014 |
|  | Polar or charged rim interface | N131, S136, R170, E190, E191, K194, N425, E427, R434, D438, S439, D444, T446, K447, D448 | N92, S97, R131, E151, E152, K155, N386, E388, R395, D399, S400, D405, T407, K408, D409 |  |
|  | Proposed hydrogen bond formed with LIMP-2 | N131, K194, R434 | N92, K155, R395 |  |
|  | Proposed salt bridge with LIMP-2 | E190, D448 | K151, D409 |  |
| Catalytic Cavity | Catalytic Residues | E274, E379 | E235, E340 | Romero et al 2019 (PDB: 1OGS, 2NSX) |
|  | Lining Residues | R159, D166, F167, W218, N273, Y283, F285, Y352, C381, S384, W420, N435, F436, V437 | R120, D127, F128, W179, N234, Y244, F246, Y313, C342, S345, W381, N396, F397, V398 |  |
| SapC Binding Interface | Domain III loop | D354, F355, L356, A357, P358 | D315, F316, L317, A318, P319 | Romero et al 2019 (PDB: 1OGS, 2NSX) |
|  | Domain III loop1-helix 6 junction | A359 | A320 |  |
|  | Domain III -helix 6 junction | K360 | K321 |  |
|  | Domain III loop 2 | K385, F386, W387, E388 | K346, F347, W348, E349 |  |
|  | Domain III loop 2-helix 7 junction | R392 | R353 |  |
|  | Domain III domain III helix 7 | D397, Q401, H404, T408 | D358, Q362, H365, T369 |  |
|  | Domain III helix 7 boundary | Y412 | Y373 |  |
|  | Domain II | K480, D482, L483, D484, R502, S503, S504, Y526 | K441, D443, L444, D445, R463, S464, S465, Y487 |  |
| Cysteine dilsulfide bridge residue | Domain I | Cys53, Cys55, Cys57, Cys62 | Cys14, Cys16, Cys18, Cys23 | Dvir et al., 2003 (PDS: 1OGS) |
| N-Linked Glycosylation Residue |  | N58, N98, N185, N309 | N19, N59, N146, N270 | Berg-Fussman et al., 1993 |

**Table 4:**
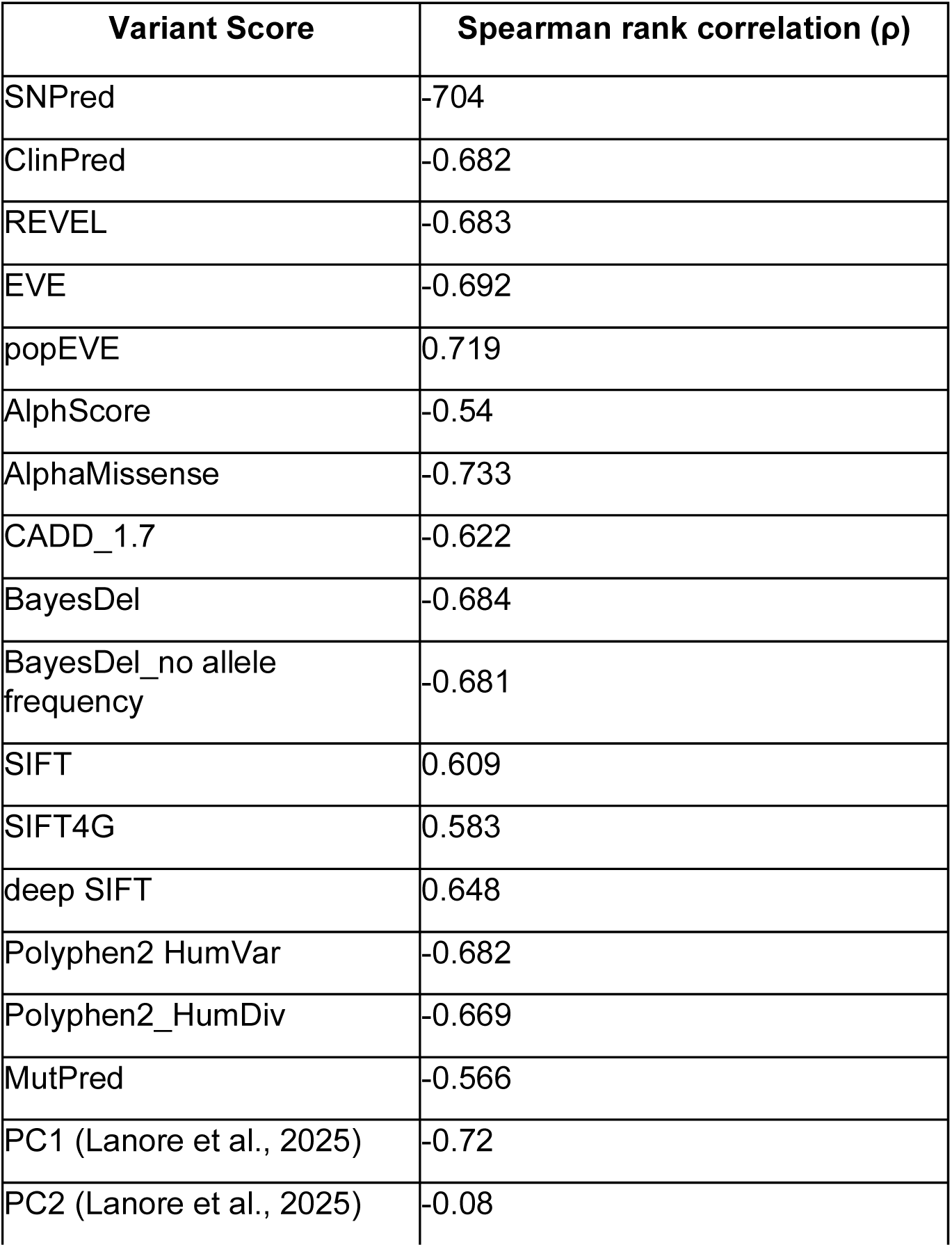
Summary of VEP correlation with *GBA1* flow-seq scores.

**Supplementary Table 5:** Correlation analyses of Flow-Seq scores with AAO and ORs across GP2 cohorts.

| Comparison | Restriction | Spearman rank (ρ) | p value | 95% CI |
|---|---|---|---|---|
| Median AAO vs Flow-Seq Score | Minimum carriers ≥ 1 | 0.244 | 3.86E-03 | [0.081, 0.395] |
|  | Minimum carriers ≥ 2 | 0.301 | 4.46E-02 | [0.008, 0.546] |
|  | Minimum carriers ≥ 5 | 0.728 | 6.12E-04 | [0.396, 0.892] |
|  | Minimum carriers ≥ 10 | 0.561 | 9.16E-02 | [-0.106, 0.880] |
| Mean AAO vs Flow-Seq Score | Minimum carriers ≥ 1 | 0.244 | 3.87E-03 | [0.081, 0.395] |
|  | Minimum carriers ≥ 2 | 0.303 | 4.34E-02 | [0.010, 0.547] |
|  | Minimum carriers ≥ 5 | 0.697 | 1.32E-03 | [0.340, 0.878] |
|  | Minimum carriers ≥ 10 | 0.539 | 1.08E-01 | [-0.137, 0.873] |
| OR vs Flow-Seq Score | Minimum allele count ≥ 5 | -0.534 | 2.71E-02 | [-0.808, -0.072] |
| OR vs Flow-Seq Score | Minimum allele count ≥ 10 | -0.473 | 1.03E-01 | [-0.812, 0.106] |

## References

Alcalay, R. N., Hsieh, F., Tengstrand, E., Padmanabhan, S., Baptista, M., Kehoe, C., Narayan, S., Boehme, A. K., & Merchant, K. (2020). Higher Urine bis(Monoacylglycerol)Phosphate Levels in LRRK2 G2019S Mutation Carriers: Implications for Therapeutic Development. Movement Disorders : Official Journal of the Movement Disorder Society, 35(1), 134–141. 10.1002/mds.27818

Alcalay, R. N., Levy, O. A., Waters, C. C., Fahn, S., Ford, B., Kuo, S.-H., Mazzoni, P., Pauciulo, M. W., Nichols, W. C., Gan-Or, Z., Rouleau, G. A., Chung, W. K., Wolf, P., Oliva, P., Keutzer, J., Marder, K., & Zhang, X. (2015). Glucocerebrosidase activity in Parkinson’s disease with and without GBA mutations. Brain : A Journal of Neurology, 138(Pt 9), 2648–2658. 10.1093/brain/awv179

Álvarez Jerez, P., Wild Crea, P., Ramos, D. M., Gustavsson, E. K., Radefeldt, M., Damianov, A., Makarious, M. B., Ojo, O. O., Billingsley, K. J., Malik, L., Daida, K., Bromberek, S., Hu, F., Schneider, Z., Surapaneni, A. L., Stadler, J., Rizig, M., Morris, H. R., Pantazis, C. B., … Blauwendraat, C. (2024). African ancestry neurodegeneration risk variant disrupts an intronic branchpoint in GBA1. Nature Structural & Molecular Biology, 31(12), 1955–1963. 10.1038/s41594-024-01423-2

Aslanzadeh, V., Brierley, G. V, Kumar, R., Çubuk, H., Vigouroux, C., Matreyek, K. A., Kudla, G., & Semple, R. K. (2025). Deep mutational scanning of the human insulin receptor ectodomain to inform precision therapy for insulin resistance. Nature Communications, 16(1), 9143. 10.1038/s41467-025-64178-4

Aslanzadeh, V., Glendinning, S., Sari, Y., Clarke, M., Rodger, C. J., Lin, X., Lenka, D. R., Richardson, L., Liang, J. R., Kleinz, T., Sproul, D., Lohmann, K., Klein, C., Sammler, E. M., Alessi, D. R., Kumar, A., Muqit, M. M. K., & Kudla, G. (2026). A Functional Genetic Atlas of Parkin Resolves Variants of Uncertain Significance and Predicts Parkinson’s Disease Age at Onset. 10.64898/2026.02.09.704817

Badonyi, M., & Marsh, J. A. (2025). <tt>acmgscaler</tt> : an R package and Colab for standardized gene-level variant effect score calibration within the ACMG/AMP framework. Bioinformatics, 41(10). 10.1093/bioinformatics/btaf503

Berg-Fussman, A., Grace, M. E., Ioannou, Y., & Grabowski, G. A. (1993). Human acid beta-glucosidase. N-glycosylation site occupancy and the effect of glycosylation on enzymatic activity. Journal of Biological Chemistry, 268(20), 14861–14866. 10.1016/S0021-9258(18)82412-7

Brady, R. O., Kanfer, J. N., & Shapiro, D. (1965). METABOLISM OF GLUCOCEREBROSIDES. II. EVIDENCE OF AN ENZYMATIC DEFICIENCY IN GAUCHER’S DISEASE. Biochemical and Biophysical Research Communications, 18, 221–225. 10.1016/0006-291x(65)90743-6

Chatterjee, D., & Krainc, D. (2023). Mechanisms of Glucocerebrosidase Dysfunction in Parkinson’s Disease. Journal of Molecular Biology, 435(12), 168023. 10.1016/j.jmb.2023.168023

Crawford, K., & Bloom, J. (2019). alignparse: A Python package for parsing complex features from high-throughput long-read sequencing. Journal of Open Source Software, 4(44), 1915. 10.21105/joss.01915

Davis, O. B., Kung, J. E., Davis, S. S., Ghosh, R., Andrews, S. V., Gould, N. S., Kluss, J. H., Thomsen, E., Agam, M., Coan, J. P., Maloney, M. T., Nguyen, A. H., Nguyen, H. N., Propson, N. E., Lozano, E. I., Yuan, T., Xa, K., Sperberg, R. A. P., Jain, S., … Henry, A. G. (2025). A Common PD-Risk GBA1 Variant Disrupts LIMP2 Interaction, Impairs Glucocerebrosidase Function, and Drives Lysosomal and Mitochondrial Dysfunction. 10.1101/2025.08.28.672891

Dekker, N., van Dussen, L., Hollak, C. E. M., Overkleeft, H., Scheij, S., Ghauharali, K., van Breemen, M. J., Ferraz, M. J., Groener, J. E. M., Maas, M., Wijburg, F. A., Speijer, D., Tylki-Szymanska, A., Mistry, P. K., Boot, R. G., & Aerts, J. M. (2011). Elevated plasma glucosylsphingosine in Gaucher disease: relation to phenotype, storage cell markers, and therapeutic response. Blood, 118(16), e118–e127. 10.1182/blood-2011-05-352971

Dobert, J. P., Schäfer, J.-H., Dal Maso, T., Ravindran, P., Huard, D. J. E., Socher, E., Schildmeyer, L. A., Lieberman, R. L., Versées, W., Moeller, A., Zunke, F., & Arnold, P. (2025). Cryo-TEM structure of β-glucocerebrosidase in complex with its transporter LIMP-2. Nature Communications, 16(1), 3074. 10.1038/s41467-025-58340-1

Dvir, H., Harel, M., McCarthy, A. A., Toker, L., Silman, I., Futerman, A. H., & Sussman, J. L. (2003). X-ray structure of human acid-β-glucosidase, the defective enzyme in Gaucher disease. The EMBO Reports, 4(7), 704–709. 10.1038/sj.embor.embor873

Galvagnion, C., Marlet, F. R., Cerri, S., Schapira, A. H. V, Blandini, F., & Di Monte, D. A. (2022). Sphingolipid changes in Parkinson L444P GBA mutation fibroblasts promote α-synuclein aggregation. Brain, 145(3), 1038–1051. 10.1093/brain/awab371

Gan-Or, Z., Amshalom, I., Kilarski, L. L., Bar-Shira, A., Gana-Weisz, M., Mirelman, A., Marder, K., Bressman, S., Giladi, N., & Orr-Urtreger, A. (2015). Differential effects of severe vs mild *GBA* mutations on Parkinson disease. Neurology, 84(9), 880–887. 10.1212/WNL.0000000000001315

García-Sanz, P., Orgaz, L., Bueno-Gil, G., Espadas, I., Rodríguez-Traver, E., Kulisevsky, J., Gutierrez, A., Dávila, J. C., González-Polo, R. A., Fuentes, J. M., Mir, P., Vicario, C., & Moratalla, R. (2017). N370S GBA1 mutation causes lysosomal cholesterol accumulation in Parkinson’s disease. Movement Disorders, 32(10), 1409–1422. 10.1002/mds.27119

Gegg, M. E., Menozzi, E., & Schapira, A. H. V. (2022). Glucocerebrosidase-associated Parkinson disease: Pathogenic mechanisms and potential drug treatments. Neurobiology of Disease, 166, 105663. 10.1016/j.nbd.2022.105663

Hassanin, E., Landoulsi, Z., Pachchek, S., Acharya, G., Aguayo, G., Alexandre, M., Ali, M., Ammerlann, W., Arena, G., Bassis, M., Batutu, R., Beaumont, K., Béchet, S., Berchem, G., Bisdorff, A., Boussaad, I., Bouvier, D., Castillo, L., Contesotto, G., … Bobbili, D. R. (2025). Penetrance of Parkinson’s disease in GBA1 carriers depends on variant severity and polygenic background. Npj Parkinson’s Disease, 11(1), 162. 10.1038/s41531-025-00997-y

Hunter, J. D. (2007). Matplotlib: A 2D Graphics Environment. Computing in Science & Engineering, 9(3), 90–95. 10.1109/MCSE.2007.55

Jack, A., Amato, D., Morris, G., & Choy, F. Y. M. (2014). Two novel mutations in glucocerebrosidase, C23W and IVS7−1 G>A, identified in Type 1 Gaucher patients heterozygous for N370S. Gene, 538(1), 84–87. 10.1016/j.gene.2014.01.015

Lanore, A., Tesson, C., Basset, A., Lejeune, F.-X., Cogan, G., Mangone, G., Sambin, S., Bertille, N., Anheim, M., Arnulf, I., Ansquer, S., Brandel, J.-P., Brefel-Courbon, C., Defebvre, L., Drapier, S., Eusebsio, A., Fabbri, M., Giordana, C., Hainque, E., … Camgrand, E. (2025). Classification of GBA1 variants and their impact on Parkinson’s disease: an in silico score analysis. Npj Parkinson’s Disease, 11(1), 226. 10.1038/s41531-025-01060-6

Lieberman, R. L. (2011). A Guided Tour of the Structural Biology of Gaucher Disease: Acid-*β*-Glucosidase and Saposin C. Enzyme Research, 2011, 1–15. 10.4061/2011/973231

Liou, B., Haffey, W. D., Greis, K. D., & Grabowski, G. A. (2014). The LIMP-2/SCARB2 Binding Motif on Acid β-Glucosidase. Journal of Biological Chemistry, 289(43), 30063–30074. 10.1074/jbc.M114.593616

Liou, B., Kazimierczuk, A., Zhang, M., Scott, C. R., Hegde, R. S., & Grabowski, G. A. (2006). Analyses of Variant Acid β-Glucosidases. Journal of Biological Chemistry, 281(7), 4242–4253. 10.1074/jbc.M511110200

Malek, N., Weil, R. S., Bresner, C., Lawton, M. A., Grosset, K. A., Tan, M., Bajaj, N., Barker, R. A., Burn, D. J., Foltynie, T., Hardy, J., Wood, N. W., Ben-Shlomo, Y., Williams, N. W., Grosset, D. G., & Morris, H. R. (2018). Features of *GBA*-associated Parkinson’s disease at presentation in the UK *Tracking Parkinson’s* study. Journal of Neurology, Neurosurgery & Psychiatry, 89(7), 702–709. 10.1136/jnnp-2017-317348

Matreyek, K. A., Stephany, J. J., & Fowler, D. M. (2017). A platform for functional assessment of large variant libraries in mammalian cells. Nucleic Acids Research, 45(11), e102–e102. 10.1093/nar/gkx183

McDonnell, A. F., Plech, M., Livesey, B. J., Gerasimavicius, L., Owen, L. J., Hall, H. N., FitzPatrick, D. R., Marsh, J. A., & Kudla, G. (2024). Deep mutational scanning quantifies DNA binding and predicts clinical outcomes of PAX6 variants. Molecular Systems Biology, 20(7), 5. 10.1038/s44320-024-00043-8

McEwen, A. E., Tejura, M., Fayer, S., Starita, L. M., & Fowler, D. M. (2026). Multiplexed assays of variant effect for clinical variant interpretation. Nature Reviews. Genetics, 27(2), 137–154. 10.1038/s41576-025-00870-x

Menozzi, E., Del Pozo, S. L., Macnaughtan, J., Mezabrovschi, R., Koletsi, S., Mitrotti, P., Gallo, L., Calabrese, R., Toffoli, M., Loefflad, N., Valzania, F., Cavallieri, F., Fioravanti, V., Yalkic, S., Limbachiya, N., Blandini, F., Avenali, M., & Schapira, A. H. (2025). Severe GBA1 variants drive the GBA1-PD clinical phenotype: implications for counselling and clinical trials. Npj Parkinson’s Disease, 11(1), 281. 10.1038/s41531-025-01063-3

Mighell, T. L., Toledano, I., & Lehner, B. (2023). SUNi mutagenesis: Scalable and uniform nicking for efficient generation of variant libraries. PLOS ONE, 18(7), e0288158. 10.1371/journal.pone.0288158

Miyoshi, K., Hagita, H., Horiguchi, T., Tanimura, A., & Noma, T. (2022). Redefining GBA gene structure unveils the ability of Cap-independent, IRES-dependent gene regulation. Communications Biology, 5(1), 639. 10.1038/s42003-022-03577-5

Nalls, M. A., Blauwendraat, C., Vallerga, C. L., Heilbron, K., Bandres-Ciga, S., Chang, D., Tan, M., Kia, D. A., Noyce, A. J., Xue, A., Bras, J., Young, E., von Coelln, R., Simón-Sánchez, J., Schulte, C., Sharma, M., Krohn, L., Pihlstrøm, L., Siitonen, A., … International Parkinson’s Disease Genomics Consortium. (2019). Identification of novel risk loci, causal insights, and heritable risk for Parkinson’s disease: a meta-analysis of genome-wide association studies. The Lancet. Neurology, 18(12), 1091–1102. 10.1016/S1474-4422(19)30320-5

Nalls, M. A., Pankratz, N., Lill, C. M., Do, C. B., Hernandez, D. G., Saad, M., DeStefano, A. L., Kara, E., Bras, J., Sharma, M., Schulte, C., Keller, M. F., Arepalli, S., Letson, C., Edsall, C., Stefansson, H., Liu, X., Pliner, H., Lee, J. H., … Singleton, A. B. (2014). Large-scale meta-analysis of genome-wide association data identifies six new risk loci for Parkinson’s disease. Nature Genetics, 46(9), 989–993. 10.1038/ng.3043

Parlar, S. C., Grenn, F. P., Kim, J. J., Baluwendraat, C., & Gan-Or, Z. (2023). Classification of GBA1 Variants in Parkinson’s Disease: The GBA1-PD Browser. Movement Disorders, 38(3), 489–495. 10.1002/mds.29314

Rizig, M., Bandres-Ciga, S., Makarious, M. B., Ojo, O. O., Crea, P. W., Abiodun, O. V., Levine, K. S., Abubakar, S. A., Achoru, C. O., Vitale, D., Adeniji, O. A., Agabi, O. P., Koretsky, M. J., Agulanna, U., Hall, D. A., Akinyemi, R. O., Xie, T., Ali, M. W., Shamim, E. A., … Global Parkinson’s Genetics Program. (2023). Identification of genetic risk loci and causal insights associated with Parkinson’s disease in African and African admixed populations: a genome-wide association study. The Lancet. Neurology, 22(11), 1015–1025. 10.1016/S1474-4422(23)00283-1

Romero, R., Ramanathan, A., Yuen, T., Bhowmik, D., Mathew, M., Munshi, L. B., Javaid, S., Bloch, M., Lizneva, D., Rahimova, A., Khan, A., Taneja, C., Kim, S.-M., Sun, L., New, M. I., Haider, S., & Zaidi, M. (2019). Mechanism of glucocerebrosidase activation and dysfunction in Gaucher disease unraveled by molecular dynamics and deep learning. Proceedings of the National Academy of Sciences, 116(11), 5086–5095. 10.1073/pnas.1818411116

Rossi, M., Schaake, S., Usnich, T., Boehm, J., Steffen, N., Schell, N., Krüger, C., Gül-Demirkale, T., Bahr, N., Kleinz, T., Madoev, H., Laabs, B. H., Gan-Or, Z., Alcalay, R. N., Lohmann, K., & Klein, C. (2025). Classification and Genotype–Phenotype Relationships of GBA1 Variants: MDSGene Systematic Review. In Movement Disorders (Vol. 40, Number 4, pp. 605–618). John Wiley and Sons Inc. 10.1002/mds.30141

Rubin, A. F., Stone, J., Bianchi, A. H., Capodanno, B. J., Da, E. Y., Dias, M., Esposito, D., Frazer, J., Fu, Y., Grindstaff, S. B., Harrington, M. R., Li, I., McEwen, A. E., Min, J. K., Moore, N., Moscatelli, O. G., Ong, J., Polunina, P. V., Rollins, J. E., … Fowler, D. M. (2025). MaveDB 2024: a curated community database with over seven million variant effects from multiplexed functional assays. Genome Biology, 26(1), 13. 10.1186/s13059-025-03476-y

Sanchez-Martinez, A., Beavan, M., Gegg, M. E., Chau, K.-Y., Whitworth, A. J., & Schapira, A. H. V. (2016). Parkinson disease-linked GBA mutation effects reversed by molecular chaperones in human cell and fly models. Scientific Reports, 6, 31380. 10.1038/srep31380

Schöndorf, D. C., Aureli, M., McAllister, F. E., Hindley, C. J., Mayer, F., Schmid, B., Sardi, S. P., Valsecchi, M., Hoffmann, S., Schwarz, L. K., Hedrich, U., Berg, D., Shihabuddin, L. S., Hu, J., Pruszak, J., Gygi, S. P., Sonnino, S., Gasser, T., & Deleidi, M. (2014). iPSC-derived neurons from GBA1-associated Parkinson’s disease patients show autophagic defects and impaired calcium homeostasis. Nature Communications, 5(1), 4028. 10.1038/ncomms5028

Shukla, N., Roelle, S. M., Suzart, V. G., Bruchez, A. M., & Matreyek, K. A. (2021). Mutants of human ACE2 differentially promote SARS-CoV and SARS-CoV-2 spike mediated infection. PLOS Pathogens, 17(7), e1009715. 10.1371/journal.ppat.1009715

Sidransky, E., Nalls, M. A., Aasly, J. O., Aharon-Peretz, J., Annesi, G., Barbosa, E. R., Bar-Shira, A., Berg, D., Bras, J., Brice, A., Chen, C.-M., Clark, L. N., Condroyer, C., De Marco, E. V., Dürr, A., Eblan, M. J., Fahn, S., Farrer, M. J., Fung, H.-C., … Ziegler, S. G. (2009). Multicenter Analysis of Glucocerebrosidase Mutations in Parkinson’s Disease. New England Journal of Medicine, 361(17), 1651–1661. 10.1056/NEJMoa0901281

Smith, L., & Schapira, A. H. V. (2022). GBA Variants and Parkinson Disease: Mechanisms and Treatments. Cells, 11(8), 1261. 10.3390/cells11081261

Sorge, J. A., West, C., Kuhl, W., Treger, L., & Beutler, E. (1987). The human glucocerebrosidase gene has two functional ATG initiator codons. American Journal of Human Genetics, 41(6), 1016–1024.

Starr, T. N., Greaney, A. J., Hilton, S. K., Ellis, D., Crawford, K. H. D., Dingens, A. S., Navarro, M. J., Bowen, J. E., Tortorici, M. A., Walls, A. C., King, N. P., Veesler, D., & Bloom, J. D. (2020). Deep Mutational Scanning of SARS-CoV-2 Receptor Binding Domain Reveals Constraints on Folding and ACE2 Binding. Cell, 182(5), 1295–1310.e20. 10.1016/j.cell.2020.08.012

Stirnemann, J., Belmatoug, N., Camou, F., Serratrice, C., Froissart, R., Caillaud, C., Levade, T., Astudillo, L., Serratrice, J., Brassier, A., Rose, C., Billette de Villemeur, T., & Berger, M. (2017). A Review of Gaucher Disease Pathophysiology, Clinical Presentation and Treatments. International Journal of Molecular Sciences, 18(2), 441. 10.3390/ijms18020441

Thomas, R., Moloney, E. B., Macbain, Z. K., Hallett, P. J., & Isacson, O. (2021). Fibroblasts from idiopathic Parkinson’s disease exhibit deficiency of lysosomal glucocerebrosidase activity associated with reduced levels of the trafficking receptor LIMP2. Molecular Brain, 14(1), 16. 10.1186/s13041-020-00712-3

Toffoli, M., Chen, X., Sedlazeck, F. J., Lee, C.-Y., Mullin, S., Higgins, A., Koletsi, S., Garcia-Segura, M. E., Sammler, E., Scholz, S. W., Schapira, A. H. V, Eberle, M. A., & Proukakis, C. (2022). Comprehensive short and long read sequencing analysis for the Gaucher and Parkinson’s disease-associated GBA gene. Communications Biology, 5(1), 670. 10.1038/s42003-022-03610-7

Tsuji, S., Choudary, P. V, Martin, B. M., Stubblefield, B. K., Mayor, J. A., Barranger, J. A., & Ginns, E. I. (1987). A mutation in the human glucocerebrosidase gene in neuronopathic Gaucher’s disease. The New England Journal of Medicine, 316(10), 570–575. 10.1056/NEJM198703053161002

Vieira, S. R. L., Mezabrovschi, R., Toffoli, M., Del Pozo, S. L., Menozzi, E., Mullin, S., Yalkic, S., Limbachiya, N., Koletsi, S., Loefflad, N., Lopez, G. J., Gan-Or, Z., Alcalay, R. N., Sidransky, E., & Schapira, A. H. V. (2024). Consensus Guidance for Genetic Counseling in *GBA1* Variants: A Focus on Parkinson’s Disease. Movement Disorders, 39(12), 2144–2154. 10.1002/mds.30006

Wang, K., Li, M., & Hakonarson, H. (2010). ANNOVAR: functional annotation of genetic variants from high-throughput sequencing data. Nucleic Acids Research, 38(16), e164–e164. 10.1093/nar/gkq603

Wang, X., Zhao, Z., Lin, L., Li, F., Qi, F., Wang, X., & Han, L. (2026). A comprehensive survey on diagnosis and assessment of Parkinson’s disease via plantar pressure analysis. Npj Parkinson’s Disease. 10.1038/s41531-026-01416-6

Waskom, M. (2021). seaborn: statistical data visualization. Journal of Open Source Software, 6(60), 3021. 10.21105/joss.03021

Wrenbeck, E. E., Klesmith, J. R., Stapleton, J. A., Adeniran, A., Tyo, K. E. J., & Whitehead, T. A. (2016). Plasmid-based one-pot saturation mutagenesis. Nature Methods, 13(11), 928–930. 10.1038/nmeth.4029

Yang, S., Gegg, M., Chau, D., & Schapira, A. (2020). Glucocerebrosidase activity, cathepsin D and monomeric α-synuclein interactions in a stem cell derived neuronal model of a PD associated GBA1 mutation. Neurobiology of Disease, 134, 104620. 10.1016/j.nbd.2019.104620

Yang, S.-Y., Beavan, M., Chau, K.-Y., Taanman, J.-W., & Schapira, A. H. V. (2017). A Human Neural Crest Stem Cell-Derived Dopaminergic Neuronal Model Recapitulates Biochemical Abnormalities in GBA1 Mutation Carriers. Stem Cell Reports, 8(3), 728–742. 10.1016/j.stemcr.2017.01.011

Yariv, B., Yariv, E., Kessel, A., Masrati, G., Chorin, A. Ben, Martz, E., Mayrose, I., Pupko, T., & Ben-Tal, N. (2023). Using evolutionary data to make sense of macromolecules with a “face-lifted” ConSurf. Protein Science, 32(3). 10.1002/pro.4582

Zhao, F., Bi, L., Wang, W., Wu, X., Li, Y., Gong, F., Lu, S., Feng, F., Qian, Z., Hu, C., Wu, Y., & Sun, Y. (2016). Mutations of glucocerebrosidase gene and susceptibility to Parkinson’s disease: An updated meta-analysis in a European population. Neuroscience, 320, 239–246. 10.1016/j.neuroscience.2016.02.007

Zhu, S., Deen, M. C., Zhu, Y., Gilormini, P., Chen, X., Davis, O. B., Chin, M. Y., Henry, A. G., & Vocadlo, D. J. (2023). A Fixable Fluorescence-Quenched Substrate for Quantitation of Lysosomal Glucocerebrosidase Activity in Both Live and Fixed Cells. Angewandte Chemie International Edition, 62(40). 10.1002/anie.202309306

